# Microbial succession in West African secondary forests: guild-specific internal stabilisation with no detectable convergence toward old-growth reference states

**DOI:** 10.64898/2026.06.19.733386

**Authors:** Anicet E. T. Ebou, Bienvenu H.K. Amani, Guy-Pacome T. Toure, Evans Ehouman, Sara Volena Zaoui, Anicet Dibonan Toure, Soda Mariane Ndiaye, Samuel Claude Yapo, Ahou Blandine Koffi, Romain Kouakou Fossou, Raphael Aussenac, Adolphe Zézé, Dominique K. Koua, Bruno Hérault

## Abstract

Secondary forest succession following agricultural abandonment is a dominant land-use transition across the tropics, yet whether soil microbial communities recover toward old-growth forest reference states remains poorly understood, particularly in West Africa. Here, we investigated the successional dynamics of bacterial and arbuscular mycorrhizal (AM) fungal communities along post-agricultural chronosequences spanning 1 to 43 years across six classified forests in Côte d’Ivoire, using Bayesian hierarchical models applied to amplicon sequencing data. Both guilds attained moderate to high alpha diversity within the first decade of succession; AM fungal diversity showed moderate evidence of age-related increase thereafter while bacterial diversity showed no directional trend. Pairwise turnover analyses revealed progressive internal convergence in AM fungal communities with plots farther apart in successional time becoming more compositionally similar, while bacterial communities showed only a weak and uncertain tendency in the same direction. Beta-dispersion modelling further indicated progressive within-forest homogenisation of AM fungal communities across abundance-weighted metrics, while bacterial assemblages showed no such stabilisation. Despite this internal convergence, compositional distances to old-growth reference plots remained persistently high for both guilds throughout the chronosequence, with no statistical evidence of recovery toward old-growth states across any dissimilarity metric or guild within the 40-year window. Indicator species analysis identified no robust stage-specific taxa after correction for multiple testing. These results indicate that microbial succession in postagricultural West African forests is characterised by rapid early reorganisation followed by stabilisation into site-specific assemblages that remain persistently distinct from old-growth reference communities. This outcome challenges the direct application of classical vegetation successional theory to soil microbiomes and indicates that no detectable recovery toward old-growth microbial community states was observed within the 40-year chronosequence.

## 1 Introduction

Secondary forest succession following agricultural abandonment is a dominant land-use transition across the tropics, playing a central role in global carbon sequestration, biodiversity recovery, and ecosystem functioning [Matsuo et al., 2021, Poorter et al., 2021]. Classical succession theory encompasses a diversity of conceptual frameworks, ranging from facilitation, tolerance, and inhibition models to non-equilibrium and state-and-transition perspectives, and encompasses both deterministic and diffuse trajectories [Suding et al., 2004]. Plant communities may thus exhibit divergent successional pathways, alternative stable states, or trajectories that remain contingent on historical and environmental context [Metzler et al., 2024]. Soil microbial communities are central to these dynamics yet are frequently overlooked in succession studies: they underpin key ecosystem processes during forest regeneration, including organic matter decomposition, nutrient cycling, and plant-soil feedbacks [Lladó et al., 2018, Ebou et al., 2024, Ran et al., 2024, Wan et al., 2024, Lian et al., 2025]. In particular, bacteria drive rapid biogeochemical transformations [Falkowski et al., 2008, Hutchins and Fu, 2017, Kuypers et al., 2018, Schäfer and Eyice, 2019], whereas arbuscular mycorrhizal (AM) fungi mediate plant nutrient acquisition and influence vegetation dynamics through obligate symbioses [Brundrett and Tedersoo, 2018, Bahadur et al., 2019, Vázquez-Santos et al., 2025]. Despite this functional importance, microbial succession has most often been inferred indirectly from vegetation recovery or soil physicochemical changes, rather than analysed as a dynamic ecological process in its own right [Fierer et al., 2010, Wetherington et al., 2022]. Furthermore, it is frequently assumed to parallel vegetation succession, implicitly adopting plant-based expectations of progressive convergence toward a forest reference state [Hart et al., 2001, Fierer et al., 2010, Knelman et al., 2012, Nemergut et al., 2013, Dini-Andreote et al., 2015, Delgado-Baquerizo et al., 2016]. Empirical evidence increasingly challenges this assumption. Microbial communities appear to respond rapidly to disturbance [Philippot et al., 2021, Liu et al., 2021], stabilise early, and remain strongly shaped by land-use legacies [Newbold et al., 2020], local environmental heterogeneity, and stochastic colonisation processes [Zhou and Ning, 2017]. High dispersal capacity, large population sizes, functional redundancy, and priority effects may further decouple microbial community trajectories from successional age [Chase, 2003, De Wit and Bouvier, 2006, Shade and Handelsman, 2012, Fukami, 2015], leading to persistent alternative community states rather than the orderly replacement through discrete stages that classical theory would predict. Yet whether these dynamics hold across the full range of tropical forest contexts remains poorly tested.

Despite growing evidence for these alternative dynamics, most empirical studies have been conducted in temperate or Neotropical forests, leaving tropical West African secondary forests severely underrepresented in the microbial succession literature. This geographical gap is consequential: West African forests face intense and ongoing anthropogenic pressure including logging, fire, and agricultural encroachment [Dago et al., 2023, Houphouet et al., 2025, Traoré et al., 2024], support high plant diversity, and harbour microbial communities structured by distinct edaphic and climatic conditions that may drive successional dynamics unlike those documented elsewhere. Secondary forests in this region show active recovery of aboveground biomass and soil organic carbon following abandonment [N’Guessan et al., 2019, Doua-Bi et al., 2021, Koffi et al., 2025], yet whether belowground microbial communities follow comparable recovery trajectories remains unknown.

Recognising these gaps, we investigate the successional dynamics of soil bacterial and AM fungal communities along post-agricultural chronosequences spanning early secondary forest to old-growth forest reference systems in Côte d’Ivoire, using an explicitly probabilistic framework that focuses on tendencies rather than deterministic endpoints [Clark, 2005, Conn et al., 2018]. We first tested whether microbial diversity trajectories conformed to a classical exponential recovery model [Hérault and Piponiot, 2018], the parametric null expected under directional convergence toward old-growth reference states, before adopting a more flexible probabilistic framework when this model proved inadequate. Rather than presuming convergence or equilibrium, we define successional properties operationally: directionality as consistent change in community composition with increasing stand age, stabilisation as declining temporal turnover or variance through time, and convergence as decreasing multivariate distance to old-growth reference communities [Anderson et al., 2006, Liu et al., 2021]. To quantify these properties while propagating uncertainty across hierarchical spatial structure and nonlinear responses, we employ Bayesian hierarchical models, which allow direct estimation of probabilistic support for alternative successional trajectories and facilitate explicit comparisons between microbial guilds [Clark, 2005]. This framework does not presume convergence or equilibrium, but instead evaluates whether and to what extent such patterns emerge from the data. By integrating temporal turnover, beta-dispersion, and distance-to-reference analyses, we aim to disentangle early reorganisation from long-term convergence and to test whether microbial succession mirrors classical vegetation-based models or instead follows alternative, non-deterministic pathways.

Specifically, we asked (i) whether microbial alpha diversity increases with successional age and differs between bacteria and AM fungi, (ii) whether microbial communities converge toward old-growth forest reference states over time, (iii) how rapidly microbial communities reorganise and stabilise following agricultural abandonment, and (iv) whether microbial succession is characterised by deterministic, stage-specific trajectories or by diffuse and probabilistic assembly processes. To address these questions, we employed four complementary analytical approaches. Alpha diversity trajectories (Q1) were characterised using Bayesian hierarchical generalised additive models, following rejection of the preliminary exponential recovery model. Compositional convergence toward old-growth reference states (Q2) was directly tested by modelling the mean dissimilarity of each secondary forest plot to old-growth plots at the same site across the chronosequence. Community reorganisation and stabilisation (Q3) were assessed through Bayesian modelling of pairwise compositional turnover and within-forest beta-dispersion. Finally, the presence or absence of stage-specific successional states (Q4) was evaluated using indicator taxa analysis.

## 2 Material and methods

### 2.1 Study sites and sampling

The study area spans a climatic and vegetation gradient across Côte d’Ivoire, with annual rainfall ranging from 1,100 mm in the north to 2,500 mm in the south. This gradient transitions from the dry, open forests of the Sudanian zone in the north, through the semi-deciduous forests of the mesophile zone in the centre, to the evergreen forests of the ombrophile zone in the south. Six classified forests were selected to represent this full gradient: Badénou and Foumbou in the northern dry zone; Haut-Sassandra and Téné in the semi-deciduous zone; and Niegré and Irobo in the evergreen zone (Figure 1).

**Figure 1:**
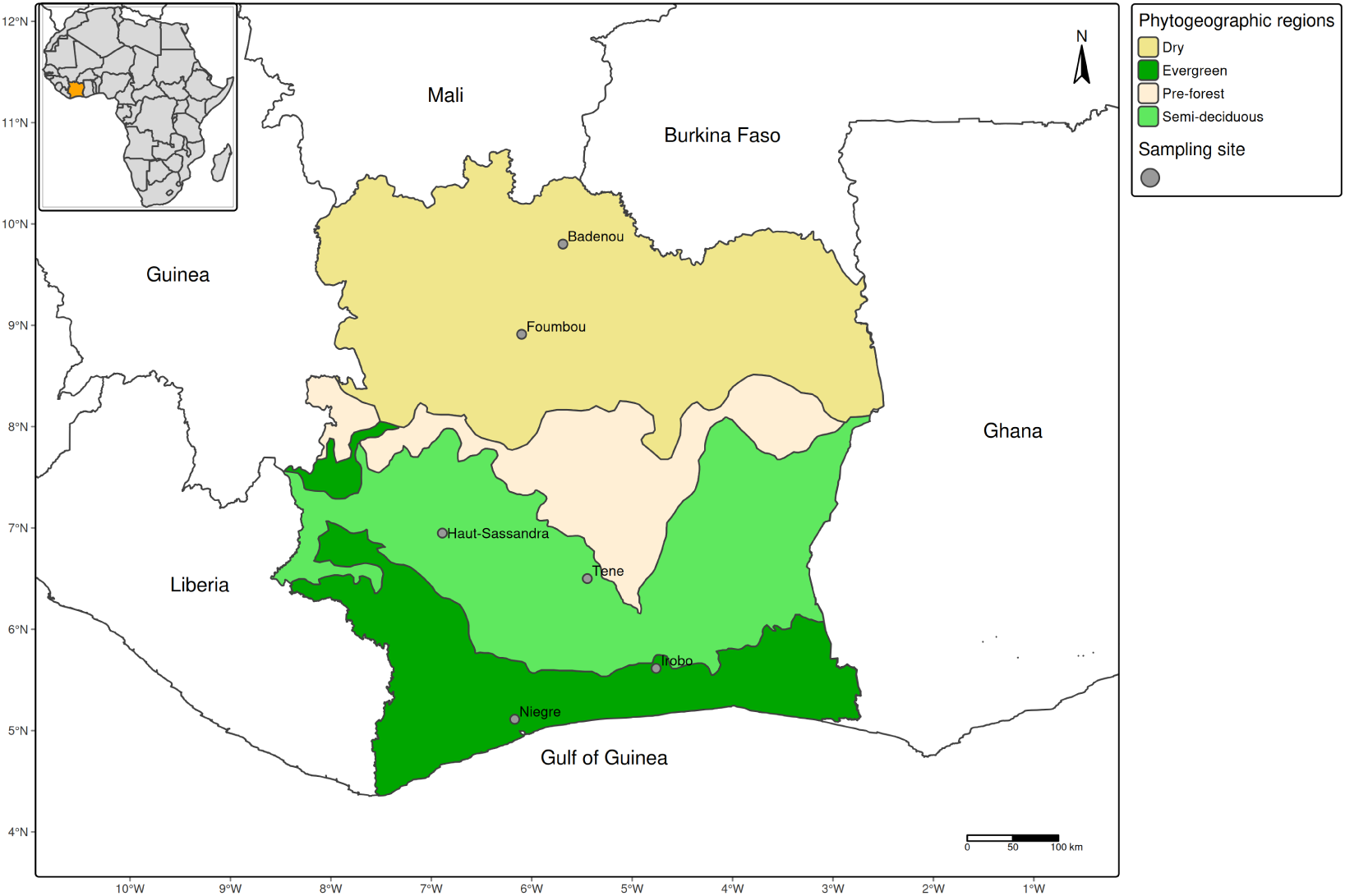
Sample sites in tropical forests across phytogeographic regions of Côte d’Ivoire (map adapted from [Guillaumet and Adjanohoun, 1971])

Plot selection followed the protocol established by Amani *et al*. (2021, 2022), using the space-for-time substitution approach, in which forests at different successional stages are used to infer ecological change over time. Forests were selected to ensure representation of the full range of phytogeographical and climatic zones of Côte d’Ivoire.

Following preliminary investigations, survey plots were randomly established within floristically homogeneous vegetation units representative of each forest. Plots representing different successional age categories were spatially distributed across each forest rather than clustered in a single location, minimising local spatial autocorrelation within sites and ensuring that the forest-level random intercept in subsequent models captures genuine between-forest heterogeneity. Plot-level metadata including exact ages, land use history, and edaphic characteristics are provided in Supplementary Table S0.

Two chronosequences were established per phytogeographical zone, for a total of six chronosequences spanning the three climatic zones of Côte d’Ivoire. Within each chronosequence, five plots of 100 m×20 m (2,000 m2) were established, comprising one old-growth forest reference plot (OGF) and four secondary forest plots spanning the following age categories since agricultural abandonment: 0-10 years (very young), 11-20 years (young), 21-30 years (mid-aged), and over 30 years (old secondary forest).

In total, 30 plots were established across the six forests (five per forest), covering a successional gradient from 1 to 43 years since agricultural abandonment, with one plot per age category per forest, yielding six plots per age category across the full dataset (0-10, 11-20, 21-30, and *>*30 years) and six old-growth forest reference plots. Plots within each forest were separated by a minimum of 150 m to reduce spatial autocorrelation.

### 2.2 Soil sampling, DNA extraction, sequencing and bioinformatics analysis

Soil sampling was conducted between September 2022 and December 2023. Each 2,000 m2 plot was divided into two 1,000 m2 subplots, with two composite samples collected per subplot; each composite consisted of four sub-samples homogenised in the field to ensure spatial representativeness. Samples were collected from the 0-10 cm mineral horizon using a soil auger, immediately placed in a portable cooler, transported to the laboratory, and stored at *−*20°C until DNA extraction. The four composite samples collected per plot were homogenised in equal proportions prior to DNA extraction, yielding a single representative soil extract per plot. All subsequent diversity and compositional analyses were therefore conducted at the plot level, with one community matrix row per plot.

DNA was extracted from 0.25 g of lysed and homogenized soil using the DNeasy PowerSoil DNA Isolation Kit (QIAGEN GmbH, Hilden, Germany) at the Laboratoire de Microbiologie, Biotechnologie et Bioinformatique, Institut National Polytechnique Félix Houphouët-Boigny (Yamoussoukro, Côte d’Ivoire). Bacterial sequences were amplified from soil DNA extracts using the 16S ribosomal RNA V3-V4 hypervariable region primers 344F (5’-ACGGRAGGCAGCAG-3’) and 802R (5’-TACCAGGGTATCTAATCCT-3’), while sequences of AM fungi were amplified using small subunit ribosomal RNA gene-specific primers WANDA (5’-CAGCCGCGGTAATTCCAGCT- 3’; [Stockinger et al., 2010]) and AML2 (5’-GAACCCAAACACTTTGGTTTCC-3’; [Lee et al., 2008]). Sequencing was performed on an Illumina MiSeq platform (GeT sequencing platform, INRAE Toulouse, France) using a 2 × 300 bp paired-end approach.

Bioinformatic analyses were conducted separately for bacterial 16S and AM fungal 18S sequences. For bacterial reads, adapters were removed with Cutadapt v2.10 [Martin, 2011], and trimmed reads were denoised with DADA2 [Callahan et al., 2016] to produce amplicon sequence variants (ASVs); taxonomic classification was performed with the Naïve Bayes RDP Classifier v2.13 [Wang and Cole, 2024]. For AM fungal reads, the gDAT pipeline [Vasar et al., 2021] was used following Vasar *et al*.(2017), with the following settings: one mismatch allowed in barcode and primer sequences, average quality threshold of 30, read pairing at 75% match threshold, chimera removal in reference mode against the MaarjAM database, and taxonomic assignment via nucleotide BLAST [Camacho et al., 2009] against MaarjAM Virtual Taxa (VT; 97% identity, 95% alignment length [Öpik et al., 2010]); reads not assigned at this step were classified against the SILVA SSU r138.2 database [Chuvochina et al., 2026].

Bacterial communities were characterised at ASV level and AM fungal communities at VT level. This asymmetry reflects a deliberate biological choice: the 18S SSU gene targeted by the WANDA/AML2 primer pair exhibits substantial intragenomic sequence variation within individual AM fungal nuclei, which can meet or exceed interspecific divergence in several Glomeromycotina lineages [Öpik et al., 2010]. Applying ASV-level resolution to AM fungal 18S amplicons would therefore yield richness estimates reflecting within-genome copy variation rather than ecological diversity among organisms. The VT framework, which clusters sequences at 97% identity against MaarjAM, was developed specifically to circumvent this problem and represents the field-established standard for AM fungal amplicon analyses [Öpik et al., 2010, Vasar et al., 2021]. A consequence of this asymmetry is that absolute diversity values are not directly comparable between guilds; all inter-guild comparisons in this study therefore concern relative patterns (directionality of diversity change, compositional turnover, beta-dispersion, and distance to reference states) computed independently within each guild. Mean sequencing depth per sample was 54 899 *±* 9 622 reads for bacteria and 11 984 *±* 8 020 reads for AM fungi.

Raw counts were retained throughout rather than rarefying to a common depth, as rarefaction introduces unnecessary variance through random subsampling and discards informative reads; the Hill-number framework and proportional-abundance dissimilarity metrics both accommodate unequal library sizes without rarefaction [McMurdie and Holmes, 2014].

### 2.3 Microbial community data and diversity metrics

Alpha and beta diversity metrics were computed from ASV and VT count tables in R v4.5.2 [R Core Team, 2025]. For each plot and guild, three alpha diversity indices were calculated using Hill-number-based metrics via the entropart package (v1.6.16; [Marcon and Hérault, 2015]): species richness, Shannon diversity, and Simpson diversity. Beta diversity was characterised using four pairwise dissimilarity metrics capturing complementary facets of compositional variation: Sørensen (presence-absence turnover), Horn and Morisita-Horn (abundance-weighted), and Bray-Curtis (abundance-weighted, sensitive to dominant taxa). Old-growth forest plots were treated as reference systems and assigned missing age values to prevent their inclusion in temporal smoothing while retaining them for comparative analyses.

### 2.4 Statistical framework

All models were fitted in a Bayesian framework using the brms package (v2.23.0; [Bürkner, 2017]), which interfaces with the probabilistic programming language Stan. Bayesian inference was adopted because it provides full posterior distributions over parameters, enables direct probabilistic statements about quantities of biological interest, and propagates uncertainty coherently across all model components. Each model was run with four Markov chain Monte Carlo (MCMC) chains of 4,000 iterations (2,000 warmup); convergence was assessed using the potential scale reduction factor (*R*^ *<* 1.01) and effective sample size diagnostics. Land use history (former crop type and years since last cultivation), soil type, and topographic position were included as fixed-effect covariates in all models to control for pre-existing site characteristics that could confound the successional age effect. Between-forest heterogeneity was accounted for through forest-level random intercepts.

### 2.5 Q1 - Alpha diversity trajectories: do bacterial and AM fungal diversity increase with successional age?

We first tested whether bacterial and AM fungal richness conformed to a classical exponential recovery model [Hérault and Piponiot, 2018, Maurent et al., 2023] as a biological validity check; results indicated that this model was inappropriate for both guilds (Supplementary Section S0), motivating adoption of a more flexible Bayesian generalised additive model (GAM) framework.

To characterise alpha diversity trajectories without imposing a parametric functional form on the age response, we fitted a Bayesian hierarchical GAM for each of the three diversity metrics (richness, Shannon, Simpson), with both guilds modelled jointly. Each model included guild-specific smooth terms over successional age, a guild main effect capturing baseline diversity differences, an old-growth forest indicator, and a forest-level random intercept:

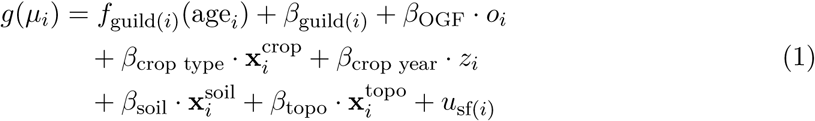

where *g*(*·*) is the link function (log); *f*_guild_(age) is a guild-specific thin-plate regression spline over successional age; 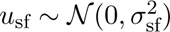 is the forest-level random intercept; and *o_i_ ∈ {*0, 1*}* is a binary indicator equal to 1 if plot *i* is an old-growth forest reference plot and 0 otherwise. The term *z_i_ ∈* R is the continuous covariate for years since last cultivation. The vectors 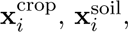 and 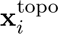 are coded indicator vectors for former crop type, soil type, and topographic position respectively, with *β*_crop type_, *β*_soil_, and *β*_topo_ denoting the corresponding vectors of contrast coefficients relative to their reference levels (*cocoa*, *lithosol*, and *peak* respectively). The term *β*_guild_(*i*) is a guild-level intercept capturing baseline diversity differences between bacteria and AM fungi.

OGF plots, for which successional age is undefined, were assigned age = NA and retained through the OGF indicator (Supplementary File S1). Species richness was modelled with a negative binomial family to account for overdispersion; Shannon and Simpson diversity were modelled with a lognormal family (Supplementary File S2).

The GAM was preferred over a linear age model for two reasons: the rejection of the exponential recovery model suggested that the age trajectory may not conform to standard parametric forms; and prior predictive checks confirmed that the thin-plate spline prior allowed both linear and nonlinear trajectories, with the data determining the degree of curvature. A sensitivity analysis comparing GAM and linear-age predictions confirmed that directional conclusions were consistent across both specifications within the 95% credible interval.

Posterior probabilities were computed from predictive distributions marginalised over forest random effects (Supplementary File S3) at representative confounder values (modal crop type, soil type, and topographic position; mean crop year). We report the probability that diversity at age 40 years exceeds diversity at age 5 years, and the probability that diversity at age 40 years meets or exceeds the predicted old-growth level.

### 2.6 Q2 - Compositional convergence toward old-growth reference states

To test Q2, we computed for each secondary forest plot the mean pairwise dissimilarity to all old-growth forest plots at the same site. This distance-to-OGF metric was modelled as a function of successional age using a Bayesian hierarchical GAM with a Beta likelihood:

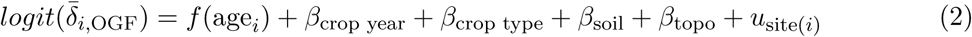

where *δ̄*_*i,*_ OGF is the mean dissimilarity of secondary forest plot *i* to all OGF reference plots at its site. A declining trajectory indicates compositional convergence toward old-growth states. The primary inferential quantity was *P* (*δ̄*_age=40_ *< δ̄*_age=5_), marginalised over site random effects and confounder values (Supplementary File S3).

### 2.7 Q3 - Community reorganisation and stabilisation: pace and guild-specificity of compositional change

Community reorganisation and stabilisation (Q3) were assessed through pairwise compositional turnover (Section 2.7.1) and beta-dispersion modelling (Section 2.7.2). Ordination and variance partitioning (Section 2.7.3) serve an exploratory role and are reported to contextualise the main results.

#### 2.7.1 Pairwise compositional turnover: primary test for directional community change with successional age

To test whether compositional dissimilarity between plot pairs increases with the age difference separating them, which is a signature of directional successional turnover, we modelled pairwise dissimilarity across all secondary forest plot combinations as a function of absolute age difference:

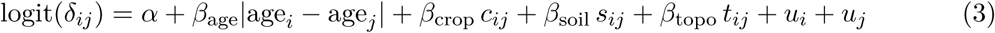

where *δ_ij_ ∈* (0, 1) denotes the pairwise dissimilarity between plots *i* and *j*, modelled using a Beta likelihood. Pair-level confounder indicators (*c_ij_, s_ij_, t_ij_ ∈ {*0, 1*}*) are binary indicators equal to 1 if plots *i* and *j* share the same former crop type, soil type, and topographic position respectively, and 0 otherwise (Supplementary File S4). They capture whether the two plots share land-use history or edaphic characteristics, thereby reducing confounding of the age-difference effect. Non-independence of pairwise data was addressed through crossed random intercepts for each plot’s forest of origin, *u_i_, u_j_ ∼ N* (0*, σ*^2^). The model was fitted independently for each combination of dissimilarity metric and microbial guild, yielding eight models per guild. To assess whether the pattern was robust to residual within-site non-independence not fully absorbed by the crossed random intercepts, a sensitivity analysis was conducted by refitting all pairwise turnover models using between-site pairs only, excluding all pairs where both plots originated from the same forest. Results of this sensitivity analysis are reported alongside the primary model results. The key quantity of interest was *P* (*β*_age_ *>* 0), the posterior probability that compositional dissimilarity increases with age difference.

#### 2.7.2 Beta-dispersion modelling: within-forest compositional stabilisation over succession

Within-forest compositional variability (beta-dispersion) was quantified as the distance of each plot to its forest-level centroid in dissimilarity space using the betadisper function from the vegan package (v2.7.2; [Oksanen et al., 2025]). These distances were modelled as a function of successional age using a Bayesian hierarchical model with a lognormal family:

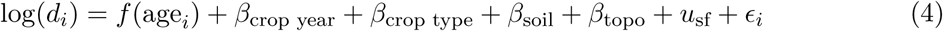

where *d_i_* is the distance of plot *i* to its forest centroid and *f* (age) is a thin-plate regression spline. Declining dispersion with age indicates progressive within-forest homogenisation (community stabilisation); increasing dispersion indicates growing within-forest variability. The primary inferential quantity was the posterior probability of dispersion declining from early (normalised age = 0.1) to late (normalised age = 0.9) succession.

#### 2.7.3 Ordination and variance partitioning: exploratory contextualisation of compositional structure (Q2–Q3)

To visualise overall patterns of compositional variation, NMDS was performed on Horn dissimilarity matrices for each guild separately using metaMDS from the vegan package (*k* = 2, up to 100 random starts). Horn dissimilarity was chosen because it accounts for species abundance while being robust to the high between-sample variation in read counts characteristic of amplicon data [Wolda, 1981]. Stress values below 0.20 were considered acceptable for a two-dimensional solution [Clarke, 1993]. To test whether successional age explained a significant proportion of compositional variation, PERMANOVA was performed using adonis2 (9,999 permutations) with forest site as a blocking factor to account for the non-independence of plots within chronosequences.

Compositional variation was formally partitioned among four predictor sets using varpart on Hellinger-transformed community matrices for secondary forest plots: **X**_1_, successional age (continuous); **X**_2_, phytogeographic zone (three levels: dry, semi-deciduous, evergreen); **X**_3_, edaphic variables (soil type and topographic position); and **X**_4_, site identity (six levels). Adjusted *R*^2^ values are reported throughout to account for differing numbers of predictors across partitions [Peres-Neto et al., 2006] (Supplementary File S5).

### 2.8 Q4 - Stage-specific versus diffuse succession: indicator taxa analysis

Stage-specific indicator taxa were identified using the IndVal association index from the indicspecies package (v1.8; [De Cáceres et al., 2010]) applied to Hellinger-transformed community tables, which reduces the influence of highly dominant taxa while preserving relative abundance structure. Associations between taxa and successional age categories were evaluated using permutation-based multilevel pattern analysis. P-values were adjusted for multiple testing using the Benjamini-Hochberg false discovery rate procedure [Benjamini and Hochberg, 1995], and only taxa retaining significance after correction were considered robust indicators.

## 3 Results

### 3.1 Q1 - Alpha diversity: early attainment and guild-specific divergence, with no progressive accumulation

Bayesian hierarchical GAMs revealed contrasting successional trajectories between guilds. Bacterial diversity showed no directional increase with age across any metric (posterior probability of increase from age 5 to 40 years: 0.15–0.26), while AM fungal diversity showed consistent moderate evidence of increase (richness: P = 0.83; Shannon: P = 0.91; Simpson: P = 0.86). Neither guild converged toward old-growth diversity levels by 40 years (bacteria: P = 0.44–0.48; AM fungi: P = 0.55–0.62). Age smooth terms did not depart substantially from linearity (*σ_sds_*: 0.325–0.414; Supplementary Table S1), and directional conclusions were consistent whether age was modelled as a smooth or linear term.

These patterns were visible in the raw data: bacterial richness spanned 576–1050 ASVs across all age categories with extensive overlap among stages, while AM fungal communities showed lower absolute diversity but similarly high within-category variability and no consistent trend with age (Figure 2).

**Figure 2:**
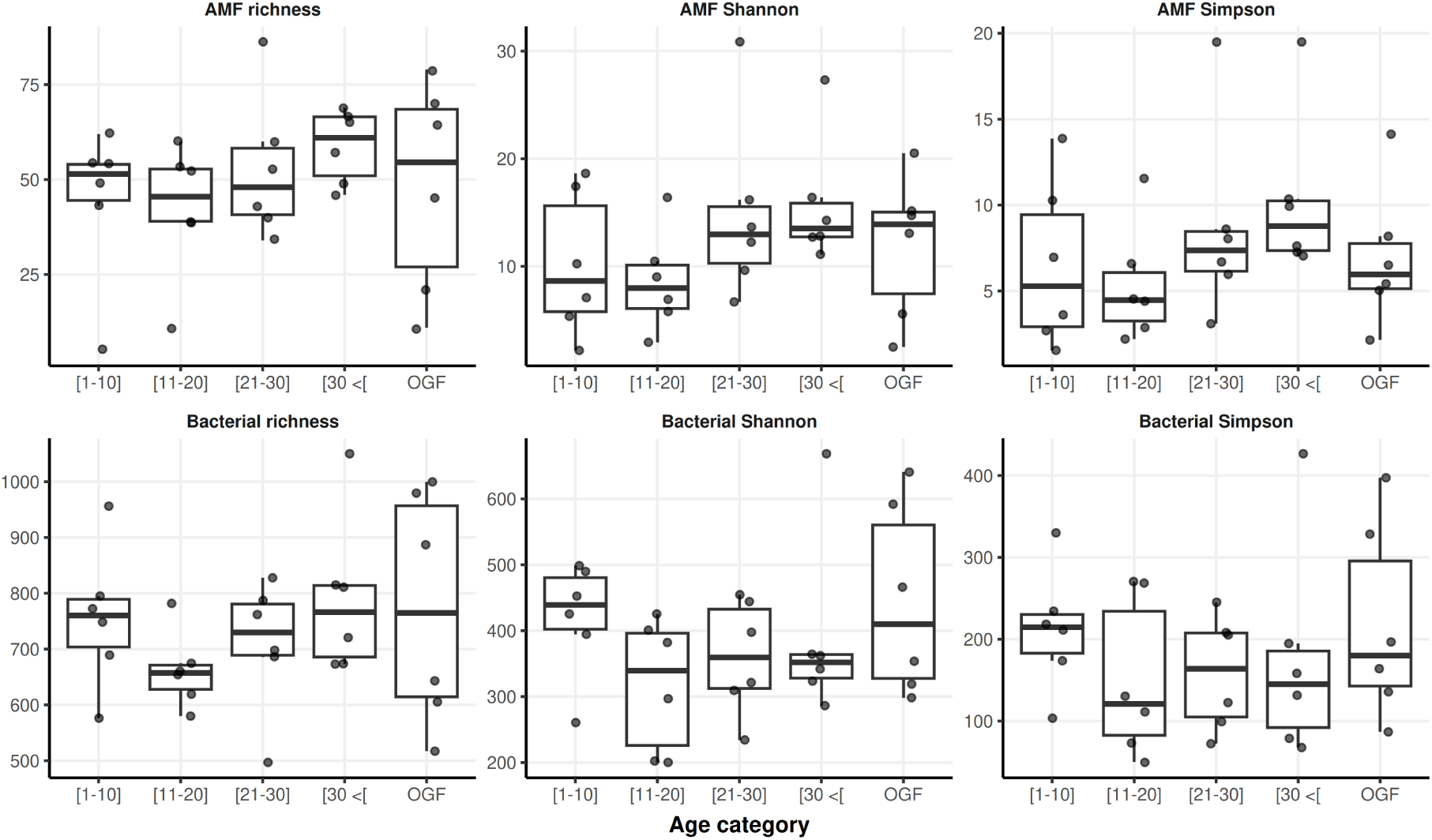
Hill-number based alpha diversity of bacterial and AM fungal communities across successional age categories in West African post-agricultural secondary forests. OGF: old-growth forest reference plots; AMF: arbuscular mycorrhizal fungi.

Bacterial communities were markedly more diverse than AM fungi in absolute terms (richness: AM fungi effect = *−*2.71, 95% CrI: [*−*2.88, *−*2.53]; Shannon: *−*3.48 [*−*3.73, *−*3.23]; Simpson: *−*3.12 [*−*3.42, *−*2.81]; Figure 3, Supplementary Tables S1–S2), though this partly reflects differences in taxonomic resolution between guilds rather than purely biological divergence. Guild-specific successional trajectories were robust to confounder inclusion: all confounder credible intervals spanned zero, and between-forest variability (forest SD: 0.28–0.31) was moderate and consistent across metrics.

**Figure 3:**
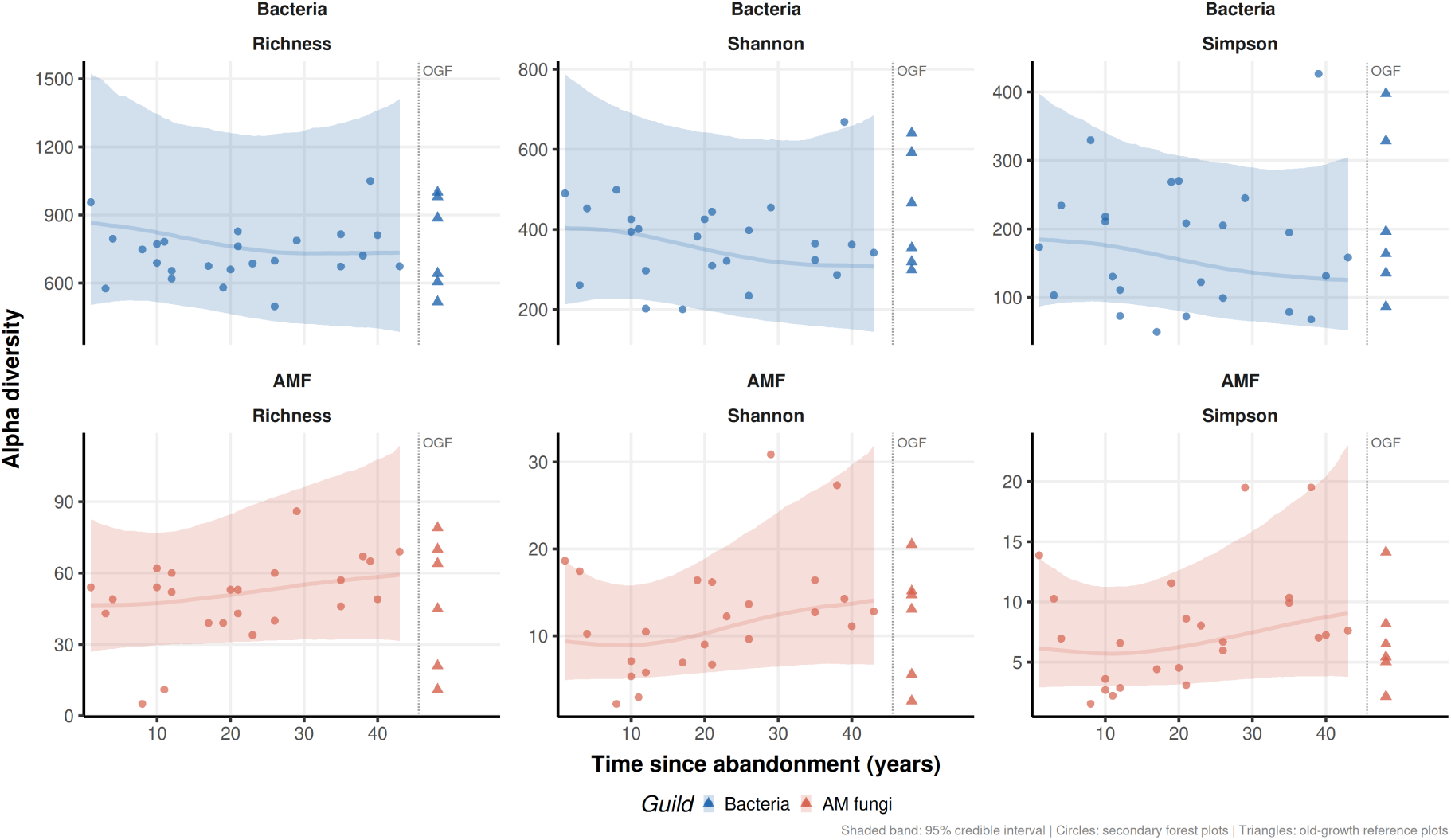
Successional dynamics of bacterial and AM fungal alpha diversity across a West African forest chronosequence. Posterior median (line) and 95% credible interval (shaded band) of predicted diversity as a function of time since agricultural abandonment. Circles represent observed values from secondary forest plots; triangles represent old-growth forest reference plots positioned beyond the dashed line. Predictions are marginalised over forest random effects and evaluated at the modal land use history and edaphic conditions across the chronosequence.

Neither guild had recovered to old-growth diversity levels by 40 years. For bacteria, posterior probabilities of reaching OGF diversity levels at age 40 were statistically indistinguishable from chance (0.43–0.48 across metrics). AM fungi showed slightly higher but still inconclusive probabilities (richness: 0.55; Shannon: 0.61; Simpson: 0.62). The old-growth forest effect was poorly estimated in all models (95% CrIs: approximately *−*2.0 to +2.0), reflecting the limited number of OGF reference plots and high between-forest heterogeneity rather than a true absence of old-growth effects.

### 3.2 Q2 - No detectable compositional convergence toward old-growth reference states within 40 years

Modelling of compositional distance to old-growth forest reference plots revealed consistently high distances for both guilds throughout the chronosequence, with no evidence of recovery toward old-growth states within the observed time frame (Figure 4, Table 1).

**Figure 4:**
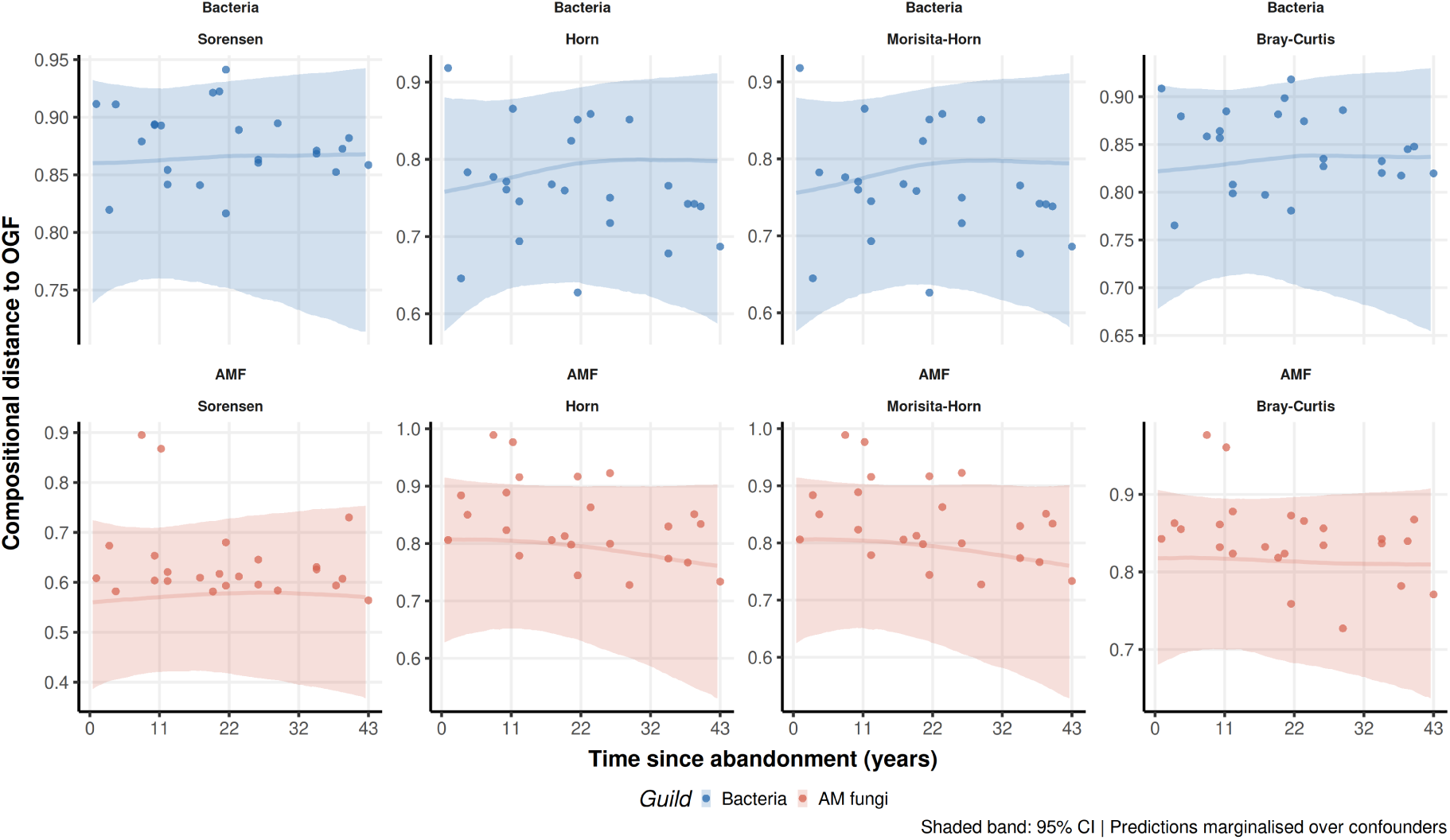
Compositional distance of secondary forest microbial communities to old-growth forest reference plots along a West African forest chronosequence. Each panel shows the posterior median predicted compositional distance (line) and 95% credible interval (shaded band) between secondary forest plots and old-growth forest reference plots as a function of time since agricultural abandonment. Points represent observed mean distances from each secondary forest plot to all old-growth forest plots of the same forest site. Predictions are marginalised over site random effects and evaluated at the modal land use history and edaphic conditions across the chronosequence. Persistent high distances across all metrics and both guilds indicate the absence of detectable compositional recovery toward old-growth reference states within 40 years of succession.

**Table 1:**
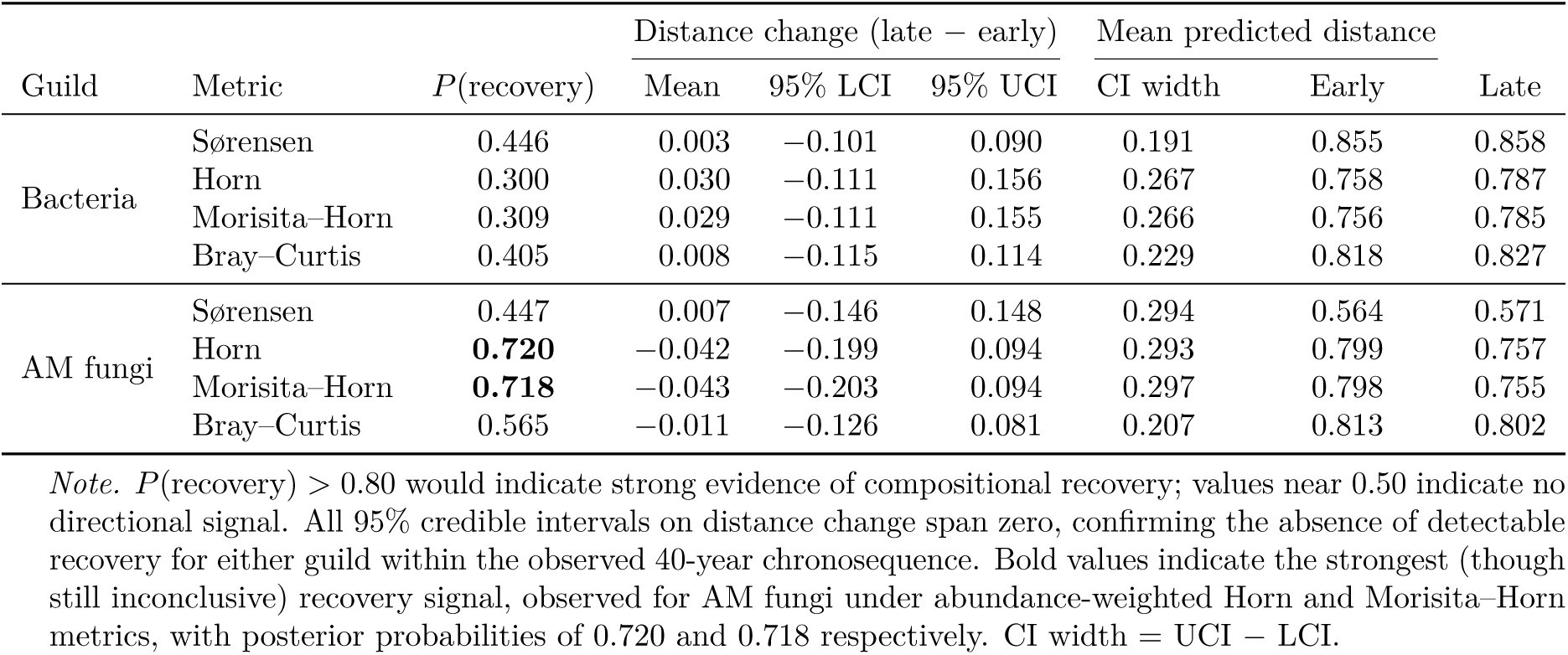
Posterior probabilities and credible intervals for compositional recovery toward old-growth forest reference states. *P* (recovery) is the posterior probability that compositional distance to old-growth forest (OGF) declines from early (normalised age = 0.1) to late (normalised age = 0.9) succession. Distance change is the posterior mean difference (late *−* early) with 95% credible interval; negative values indicate declining distance (recovery). CI width is the width of the 95% credible interval on distance change. Predictions are marginalised over site random effects and evaluated at modal confounder values.

Compositional distances to old-growth reference plots remained persistently high for both guilds throughout the chronosequence, with no evidence of recovery toward old-growth states within 40 years. Bacterial distances were high and stable across all metrics (early succession: 0.76–0.86; late succession: 0.79–0.86), with posterior probabilities of recovery near chance (Sørensen: P = 0.45; Horn: P = 0.30; Morisita-Horn: P = 0.31; Bray-Curtis: P = 0.41) and all credible intervals spanning zero. AM fungal distances were similarly high and persistent (early: 0.56–0.81; late: 0.76–0.80), with marginally stronger but still inconclusive evidence for recovery on abundance-weighted metrics (Horn: P = 0.72; Morisita-Horn: P = 0.72) and credible intervals spanning zero across all metrics.

An important caveat applies: the six OGF reference plots span three phytogeographic zones and are themselves highly compositionally heterogeneous (mean pairwise OGF distances: 0.697–0.900 across metrics and guilds; Supplementary Table S5), comparable to within-secondary forest variation. Within-SF dispersion remained substantial (bacteria: 0.352–0.478; AM fungi: 0.356–0.478), confirming that the high distances between secondary and old-growth communities primarily reflect genuine dissimilarity rather than artefactual inflation by OGF heterogeneity. Nevertheless, the limitation of a single OGF reference plot per forest site introduces irreducible uncertainty into landscape-scale recovery assessments.

### 3.3 Q3 - Rapid early reorganisation followed by internal stabilisation, with guild-specific dynamics

#### 3.3.1 Pairwise turnover: progressive compositional convergence in AM fungi, not bacteria

Pairwise compositional turnover analyses revealed a counterintuitive pattern: compositional dissimilarity between plot pairs decreased, rather than increased, with age difference across all four dissimilarity metrics and both guilds (Supplementary Table S3). For AM fungi, posterior probabilities of dissimilarity increasing with age difference were consistently low (Sørensen: P = 0.07; Horn: P = 0.07; Morisita-Horn: P = 0.06; Bray-Curtis: P = 0.04), with negative age-difference effects across all metrics (range: -0.322 (Horn) to -0.205 (Sørensen)). This result indicates that plots farther apart in successional time are compositionally more similar, which is consistent with progressive internal convergence toward a shared late-successional composition. Bacterial communities showed a weaker and more uncertain tendency in the same direction (age effect range: *−*0.14 to *−*0.19; P range: 0.09–0.23). Although this analysis generates 276 pairs from 24 secondary forest plots, the effective number of independent observations remains constrained by the underlying 24 plots and results should be interpreted accordingly.

#### 3.3.2 Beta-dispersion: within-forest homogenisation in AM fungi only

Beta-dispersion modelling further supported guild-specific differences in community stabilisation (Supplementary Figure S1, Supplementary Table S4). Bacterial dispersion showed no evidence of decline across any dissimilarity metric (P range: 0.17–0.31), with positive dispersion change estimates indicating a tendency toward increasing rather than decreasing within-forest variability. AM fungal dispersion showed moderate to strong evidence of decline for abundance-based metrics (Horn: P = 0.73; Morisita-Horn: P = 0.75; Bray-Curtis: P = 0.73), with weaker evidence for Sørensen dissimilarity (P = 0.50). Together, declining pairwise turnover and declining dispersion in AM fungi indicate progressive internal stabilisation, while bacteria show neither signal.

#### 3.3.3 Contextualising evidence: forest identity, not successional age, organises compositional space

NMDS ordinations based on Horn dissimilarities yielded acceptable two-dimensional solutions (bacteria: stress = 0.17; AM fungi: stress = 0.18). Plots from different successional stages broadly overlapped in ordination space for both guilds, with no directional structuring along the chronosequence; forest identity was a visually stronger organising factor than successional age (Figure 5). PERMANOVA confirmed that successional age explained only a small and non-significant proportion of compositional variation (bacteria: pseudo-*F* = 1.106, Adjusted *R*^2^ = 0.002, *P* = 0.212; AM fungi: pseudo-*F* = 0.996, Adjusted *R*^2^ = 0.0001, *P* = 0.440; 9,999 permutations), with forest site accounting for a substantially larger fraction of variance in both guilds.

**Figure 5:**
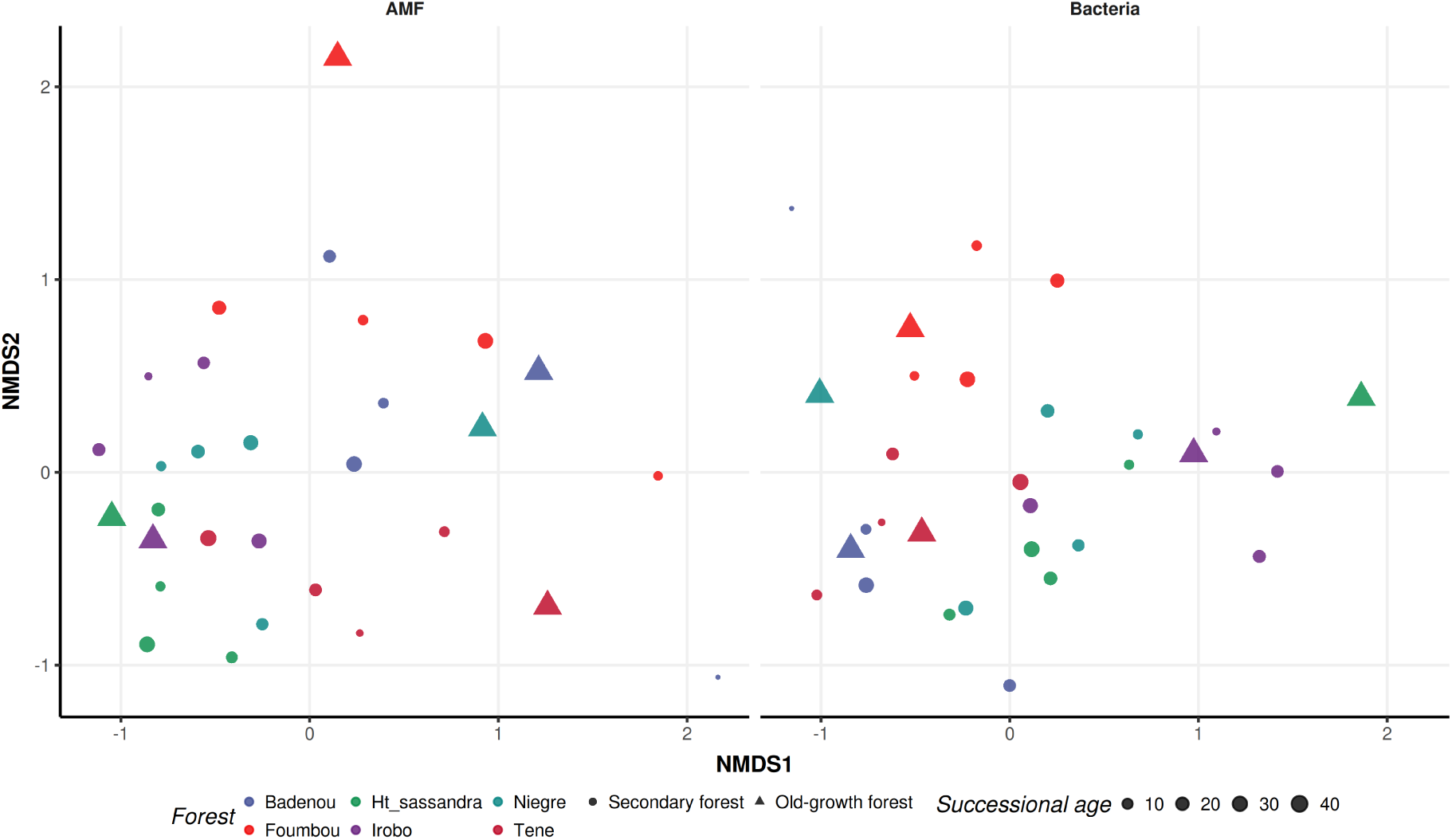
Non-metric multidimensional scaling ordination of bacterial and AM fungal community composition across a West African forest chronosequence. Two-dimensional NMDS solutions based on Horn dissimilarities for AM fungal (left) and bacterial (right) assemblages. Each point represents one forest plot, coloured by forest of origin; point size is scaled proportionally to successional age for secondary forest plots (circles), while old-growth forest plots (triangles) are shown at a fixed large size to distinguish them as a structurally distinct category without implying a numerical age.

Variance partitioning revealed strong collinearity among the four predictor sets, precluding meaningful interpretation of individual partitioned fractions (full results in Supplementary Section S6). Across both guilds, successional age explained a negligible to negative unique fraction of compositional variation when conditioned on the remaining predictors, confirming the primacy of site identity and phytogeographic context over successional time.

### 3.4 Q4 - Diffuse assembly rather than discrete successional stages

Indicator species analysis provided no support for discrete, stage-specific successional states in either guild. Of 11,857 bacterial taxa, 65 showed nominal IndVal associations prior to FDR correction, the majority with the 1–10 year category (29 taxa; typically A *≥* 0.80, B = 0.40–0.80), with smaller numbers associated with intermediate (5 taxa), late secondary (13 taxa), and old-growth categories (6 taxa). None retained significance after correction (all adjusted P = 1). No AM fungal taxa showed significant associations at any threshold before correction. However, with only 220 AM fungal taxa and a maximum of one plot per design cell, the FDR correction could eliminate all associations regardless of any true biological signal; the AM fungal null result should therefore be interpreted as uninformative rather than as evidence against stage-specific taxa.

## 4 Discussion

Across six West African forest chronosequences and four complementary analytical dimensions (alpha diversity, compositional distance to old-growth references, beta-dispersion, and pairwise compositional turnover), a coherent picture of microbial successional dynamics emerges: rapid early community assembly followed by prolonged stabilisation, with successional age exerting only weak and uncertain control over community structure. Before discussing these patterns, two caveats deserve emphasis. First, with 30 plots distributed across five successional age categories and six sites spanning three phytogeographic zones, many design cells contain a single plot; the analyses should therefore be interpreted as hypothesis-generating rather than confirmatory, and the Bayesian framework is used precisely because it represents this uncertainty honestly. Second, the observed divergence between guilds, which challenges the assumption that bacteria and AM fungi track a common successional trajectory, may partly reflect methodological asymmetries between ASV-level and VT taxon-level resolution, rather than purely biological processes; inter-guild comparisons should be read with this caveat in mind.

A third caveat concerns the space-for-time substitution design itself: by inferring temporal trajectories from plots at different successional stages rather than tracking individual plots longitudinally, the chronosequence approach assumes that plots differing in age are otherwise comparable. However, such assumption that the strong site identity effects documented throughout this study suggest, is only partially met. Longitudinal monitoring of the same plots over time would be necessary to confirm whether the patterns described here reflects genuine within-plot successional trajectories.

### 4.1 Alpha diversity is not a reliable indicator of successional trajectory

Alpha diversity provides limited information on successional trajectory in this system: both guilds attained moderate to high diversity within the first decade of abandonment, and neither showed the progressive accumulation predicted by classical succession theory. This early stabilisation, rather than any directional trend, is the dominant signal across all three indices and both guilds.

Bacterial communities in the 1–10 year category already displayed high richness (576–956 ASVs), with Hill-based Shannon and Simpson diversity remaining comparably high and broadly variable across all subsequent age categories including old-growth forest. AM fungal communities showed substantially lower absolute diversity than bacteria — a difference confirmed across all three indices and all Bayesian models with large and precisely estimated guild effects (richness: AMF effect = *−*2.71, 95% CrI: [*−*2.88, *−*2.54]) — but similarly exhibited high within-category variability and no consistent increase with age.

Bayesian hierarchical models reinforced these patterns quantitatively. The posterior probability of bacterial diversity increasing from age 5 to 40 years ranged from only 0.15 to 0.26 across all indices, values below or near chance that indicates no directional increase, while AM fungi showed moderate but consistent evidence of increase (richness: P = 0.83; Shannon: P = 0.91; Simpson: P = 0.86). Neither guild showed clear convergence toward old-growth diversity levels by 40 years, with posterior probabilities of OGF-level recovery near chance for bacteria (0.44-0.48) and inconclusive for AM fungi (0.55-0.62).

The absence of progressively increasing richness or effective diversity indicates that microbial communities rapidly reassemble a large fraction of their local species pool early in succession, rather than accumulating diversity gradually with forest age [Nemergut et al., 2013, Ning et al., 2020]. For bacteria, this early stabilisation is consistent with high dispersal capacity, large population sizes, and functional redundancy, which facilitate rapid recolonisation once basic soil conditions are restored following agricultural disturbance [Finlay, 2002, Zhou and Ning, 2017, Ning et al., 2019, Lennon and Jones, 2011, Anthony et al., 2023, Bradley, 2025].

The rapid early establishment of bacterial communities may also reflect their central role in decomposing the accumulating woody debris and leaf litter characteristic of early secondary succession, which drives nutrient release and creates positive feedbacks for community assembly [Hérault et al., 2010]. For AM fungi, the pronounced within-category variability and moderate but uncertain age-related increase suggest that diversity is more strongly shaped by local host plant composition, microsite heterogeneity, and land-use legacies than by successional age alone [Davison et al., 2015, Brundrett and Tedersoo, 2018, Zhou et al., 2018]. The robustness of these patterns to confounder inclusion, crop type, crop year, and edaphic variables all showing credible intervals spanning zero, indicates that they reflect genuine features of the microbial successional process rather than artefacts of differential land use legacy across age classes. Taken together, the extensive overlap of diversity metrics across age categories confirms that alpha diversity is a poor indicator of longer-term successional trajectory or ecosystem recovery in this system [Shade and Handelsman, 2012, Cheng et al., 2022, Ramond et al., 2025].

### 4.2 No convergence toward old-growth forest states within 40 years

Neither bacterial nor AM fungal communities showed detectable convergence toward oldgrowth forest compositional states across 40 years of succession, a result that held across all four dissimilarity metrics and both guilds. This finding must however be contextualised by a critical feature of the reference design: the six OGF plots span three phytogeographic zones and are themselves highly compositionally heterogeneous at the landscape scale, with mean pairwise OGF distances (0.697-0.900) comparable to within-secondary forest variation. The OGF assemblages therefore do not constitute a single coherent successional target, as a secondary forest in the evergreen south cannot biologically be expected to converge toward an OGF community from the dry northern zone, and the non-convergence result cannot be cleanly separated from this reference heterogeneity without a denser OGF sampling design.

Compositional distances to OGF reference plots remained persistently high for both guilds throughout the chronosequence (bacteria: 0.76–0.86 at early succession, 0.79–0.86 at late succession; AM fungi: 0.56-0.81 at early succession, 0.76-0.80 at late succession). Posterior probabilities of recovery were near or below chance for bacteria across all metrics (Sørensen: P = 0.45; Horn: P = 0.30; Morisita-Horn: P = 0.31; Bray-Curtis: P = 0.41) and only marginally stronger for AM fungi on abundance-weighted metrics (Horn: P = 0.72; Morisita-Horn: P = 0.72), with all credible intervals on distance change spanning zero. Secondary forest microbial communities are thus stabilising into compositional states that are internally consistent but persistently distinct from old-growth references, a pattern more consistent with historically contingent assembly than with directional recovery toward a forest climax [Prosser et al., 2007, Oehl et al., 2003, Mori et al., 2018, Lu et al., 2019].

The broader structure of compositional variation reinforces this picture. In NMDS ordination space, plots from different successional stages broadly overlap, confirming that successional stage alone does not impose detectable shifts in community centroids. Forest identity emerged as a consistently stronger organising factor than successional age, reflecting substantial between-site heterogeneity that persisted even after accounting for land use history and edaphic variables in hierarchical models (Adj. *R*^2^: 0.084), consistent with strong site-level filtering mediated by soil physical structure, local plant communities, and historical contingency [Wu et al., 2017, Wang et al., 2019, Guasconi et al., 2023, Zhang et al., 2024].

The posterior probabilities of OGF-level recovery (0.43–0.62 across metrics and guilds) should be read as expressions of genuine uncertainty rather than as evidence of partial recovery or its absence. This uncertainty stems from two inseparable sources: the limited number of OGF reference plots per site, and the cross-zone compositional heterogeneity among OGF plots that prevents them from constituting a single biologically coherent target. We make the modest claim that within the 40-year window observed, secondary forest microbial communities show no detectable trajectory toward old-growth compositional states and that site identity organises community composition more strongly than successional age. We do not argue that alternative stable states exist in the formal theoretical sense, which would require evidence of resistance to perturbation and multi-stability beyond what a single-timepoint chronosequence can provide. Critically, community stabilisation and community recovery are decoupled processes: the former can occur robustly without the latter, as the following section demonstrates.

### 4.3 Rapid reorganisation followed by internal stabilisation, with guild-specific dynamics

Both guilds reorganise compositionally within the first years after agricultural abandonment and subsequently stabilise into site-specific assemblages, a process that is guild-specific in both pace and magnitude and consistently more pronounced in AM fungi than in bacteria.

Pairwise turnover analyses revealed a consistent and counterintuitive pattern: across all four dissimilarity metrics and both guilds, compositional dissimilarity between plot pairs decreased rather than increased with age difference. For AM fungi, posterior probabilities of dissimilarity increasing with age difference were consistently low (Sørensen: P = 0.12; Horn: P = 0.09; Morisita-Horn: P = 0.08; Bray-Curtis: P = 0.07), with negative age-difference effects across all metrics (range: *−*0.20 to *−*0.36). Bacterial communities showed the same tendency, though considerably more weakly (P range: 0.14–0.29; effect range: *−*0.11 to *−*0.16). Plots farther apart in successional time are thus compositionally more similar to one another which is a pattern inconsistent with ongoing directional turnover toward a diverging climax, and more consistent with progressive internal convergence toward a shared late-successional composition [Allison and Martiny, 2008, Chase, 2010, Vellend, 2010, Chase and Myers, 2011, Delgado-Baquerizo et al., 2016, Mori et al., 2018]. This convergence was more pronounced for AM fungi than for bacteria, consistent with the stronger directional signal in AM fungal alpha diversity and dispersion analyses.

Three alternative explanations for this negative age-difference effect deserve consideration. First, and arguably most parsimonious, early-successional communities may already encompass much of the regional species pool through rapid recolonisation, such that both young and old forests share a broadly similar set of taxa from the outset; under this scenario, the negative age-difference effect would reflect early attainment of a shared regional pool rather than progressive convergence over time [Nemergut et al., 2013, Fukami, 2015, Shade and Handelsman, 2012, Peay et al., 2016, Vannette and Fukami, 2014]. This interpretation is consistent with the early stabilisation of alpha diversity in both guilds and the absence of directional age effects in bacterial communities. The current chronosequence design cannot adjudicate between these mechanisms, as its minimum observed age of approximately one year does not capture the immediate post-disturbance period; plots from the first months following abandonment, combined with a null model of random assembly from the regional pool, would be required to do so. Second, the crossed random intercepts in the pairwise model may not fully absorb the non-independence of pairs sharing a forest site [Harmon and Glor, 2010, Lichstein, 2007, Anderson et al., 2011, Zelený, 2018]. A sensitivity analysis restricted to between-site pairs addressed this concern: negative age-difference effects were maintained across all metrics and both guilds, with AM fungal posterior probabilities of increase remaining low (Sørensen: P = 0.15; Horn: P = 0.13; Morisita-Horn: P = 0.13; Bray-Curtis: P = 0.10) and effect sizes of similar magnitude (range: *−*0.20 to *−*0.32), ruling out within-site non-independence as a driver of the pattern. Third, the negative age-difference effect for bacteria does not constitute strong evidence under any model specification and is reported for completeness only; the convergence signal is specific to AM fungal communities and should not be generalised to bacteria without further evidence [Clark, 2005, Conn et al., 2018, Dini-Andreote et al., 2015, Zhou and Ning, 2017].

The apparent tension between the bacterial alpha diversity result, no directional trend, with posterior probabilities of increase below chance (0.15–0.26), and the weak negative age-difference effect in pairwise turnover (P range: 0.09–0.23) deserves explicit reconciliation. These findings are not contradictory: a community can change in species composition without changing in richness or effective diversity if taxa are replaced by others of similar diversity but different identity, leaving alpha diversity invariant while generating beta diversity signals. In bacteria, however, the turnover signal is too weak and uncertain (P values approaching 0.5 under several metrics) to be interpreted as evidence of directional change. The overall picture for this guild is one of compositional stasis following rapid early reorganisation: no systematic accumulation of diversity with age, no progressive within-forest homogenisation, no convergence toward OGF states, and no robust indicator taxa.

Beta-dispersion modelling reinforced this guild contrast. Bacterial within-forest compositional variability showed no evidence of decline across any dissimilarity metric (P range: 0.14–0.22), with positive dispersion change estimates indicating a tendency toward increasing rather than decreasing heterogeneity with age. In contrast, AM fungal dispersion declined with moderate to strong evidence for all abundance-weighted metrics (Horn: P = 0.74; Morisita-Horn: P = 0.75; Bray-Curtis: P = 0.73), indicating progressive within-forest homogenisation over succession; the weaker evidence for Sørensen dissimilarity (P = 0.56) suggests this homogenisation operates primarily at the level of abundance structure rather than species presence-absence. Together, declining turnover and declining dispersion in AM fungi point to a process of internal stabilisation that mirrors patterns described for plant communities and ectomycorrhizal fungi in temperate successional systems [Dumbrell et al., 2011, Zhou and Ning, 2017], here documented for the first time in a tropical post-agricultural context. That bacteria show none of these signals while AM fungi show all of them consistently across metrics likely reflects the tighter coupling of AM fungi to shifting plant communities and the greater independence of bacterial assembly from vegetation dynamics. This guild contrast is consistent with the broader finding that mycorrhizal symbiosis types are strongly structured by climate and vegetation context at global scales [Steidinger et al., 2019], suggesting that the successional dynamics of AM fungal communities in West African forests may be particularly sensitive to the progressive change in plant community composition along the chronosequence.

### 4.4 Succession is diffuse and probabilistic, not organised around discrete stages

Indicator taxa analysis provided no support for discrete, stage-specific successional states in either guild, confirming that microbial succession in this system is not organised around orderly community replacement.

Of 11,857 bacterial taxa, 65 showed nominal IndVal associations with successional categories prior to correction, the majority with early succession (1–10 years; 29 taxa), with smaller numbers associated with intermediate and late stages, but none retained significance after false discovery rate correction. No AM fungal taxa showed significant associations with any successional category at any threshold. For bacteria, the absence of robust indicator taxa after FDR correction, despite 65 nominal associations prior to correction, indicates that succession proceeds through diffuse, probabilistic shifts in relative abundance among broadly distributed taxa rather than through discrete sets of stage-specific specialists. [Davison et al., 2015, Lu et al., 2019]. For AM fungi, the null result should be interpreted cautiously: with only 220 taxa and a maximum of one plot per design cell, the FDR procedure could eliminate all associations regardless of any true biological signal, and the absence of indicator taxa is therefore uninformative rather than evidence against stage-specific assembly. This caveat notwithstanding, the absence of detectable plant-fungal successional coupling is particularly striking given that AM fungi are obligate plant symbionts for which tighter coupling might have been expected. Its absence suggests that host availability alone is insufficient to determine AM fungal community composition, and that dispersal limitation, priority effects, and soil legacy effects introduce substantial stochasticity into AM fungal assembly even when host communities are changing directionally.

Taken together, these results support the conclusion that microbial succession in post-agricultural West African forests follows a diffuse, contingent trajectory rather than the orderly stage-replacement implied by classical vegetation succession models. This conclusion is robust across both guilds, all analytical dimensions, and the full 40-year span of the chronosequence. The balance between stochastic and deterministic processes in community assembly has long been recognised as a central axis of variation in ecology [Hérault, 2007], and the present results suggest that stochastic processes dominate microbial community assembly in this system, at least over the observed 40-year window.

These findings carry direct implications for forest restoration practice in West Africa and comparable tropical post-agricultural landscapes. The decoupling of internal microbial stabilisation from compositional recovery toward old-growth states means that commonly used indicators of restoration success, stand age, canopy cover, plant species richness, may be poor proxies for belowground microbial recovery. Practitioners targeting microbially mediated ecosystem functions (carbon sequestration, nitrogen cycling, mycorrhizal plant support) should not assume passive recovery in parallel with aboveground vegetation. The stronger directional signals in AM fungi relative to bacteria further suggest that plant–fungal symbiotic networks may be a more tractable target for active intervention than the broader bacterial community, which reorganises rapidly post-disturbance but then stabilises into a state persistently distinct from old-growth regardless of successional age.

## 5 Conclusion

Across six West African post-agricultural forest chronosequences and four complementary analytical frameworks, the evidence consistently points to rapid early microbial community reorganisation followed by progressive internal stabilisation into site-specific assemblages that remain compositionally distinct from old-growth forest references over 40 years of succession. This process is clear and consistent for AM fungal communities across all beta diversity dimensions, but weak or absent for bacteria. Rather than mirroring the progressive, directional convergence predicted by classical vegetation succession theory, microbial communities, particularly bacteria, show no consistent successional trajectory in alpha diversity or compositional space. AM fungal communities display clearer signatures of progressive internal homogenisation, yet remain compositionally distant from old-growth reference states throughout the chronosequence, and no guild produced robust stage-specific indicator taxa. These findings collectively indicate that microbial succession is diffuse and probabilistic, with site identity explaining more compositional variation than successional age. However, the relative contribution of stochastic versus deterministic assembly processes, while consistent with a stochasticity-dominated system, was not formally quantified with null-model approaches and remains a hypothesis to be tested in future work. The decoupling of microbial and vegetation recovery trajectories documented here underscores the need to monitor belowground communities explicitly in forest restoration programmes. Whether active intervention strategies, such as soil inoculation from old-growth reference plots or management to reduce AM fungal dispersal barries, could accelerate belowground recovery remains an open and empirically testable question. These approaches are proposed as future research priorities rather than direct recommendations from the present results. Moreover, longer chronosequences, longitudinal monitoring, and denser old-growth reference sampling will be essential to determine whether the compositional gap between secondary and old-growth microbial communities eventually closes, and on what timescales.

## Acknowledgments

This work was done under the framework of the MIDAS Project that was financially supported by the British Ecological Society Ecologist in Africa grant number EA22/1326 and the TERRI4SOL project (CIRAD), which benefited from a grant from the Fonds Français pour l’Environnement Mondial (FFEM) under the Agence Française de Développement (AFD) grant agreement number CCI 1711.01 D.

## Competing interests

The author(s) declare no competing interests.

## Data Availability

Detailed material and methods along with model parameters and posteriors are available in the supplementary material file.

Raw sequencing reads are available on the National Center for Biotechnological Information (NCBI) under project ID PRJNA1440042.

Information on software and reproducibility is available in Supplementary File S6.

The code and data that support the findings of this study are openly available on Zenodo at https://doi.org/10.5281/zenodo.20931884 and GitHub at https://github.com/Ebedthan/midas_microbial_succession_code.

## Notes

https://github.com/Ebedthan/midas_microbial_succession_code

https://doi.org/10.5281/zenodo.20931884

### Competing Interest Statement

The authors have declared no competing interest.

### Summary of Updates

This version of the manuscript has been revised to update the title, the material and methods and the discussion along with an urgent fix in PERMANOVA result interpretation. All these change was made to better fit the study results and to clarify and make the manuscript consistent for the reader

