## Supplementary material for "Microbial succession in West African secondary forests: guild-specific internal stabilisation with no detectable convergence toward old-growth reference states"

**Supplementary Table S0. Plot-level metadata for all 30 survey plots across six classified forests in Côte d'Ivoire.**

Table 1: Supplementary Table S0. Plot-level metadata for all 30 survey plots across six classified forests in Côte d'Ivoire. Age: time since agricultural abandonment in years; OGF = old-growth forest reference plot (age undefined). Age category: successional stage class. Phyto: phytogeographic zone. Crop type: former agricultural crop (not applicable for OGF plots). Crop year: years since last cultivation at time of sampling (not applicable for OGF plots). Soil type and topographic position follow the classification used in all statistical models; reference levels used in Bayesian models are lithosol (soil type) and summit (topography). Coordinates are in decimal degrees (WGS84).

| **Plot** | **Forest** | **Phyto. zone** | **Age (yr)** | **Latitude** | **Longitude** | **Age category** | **Crop type** | **Crop year** | **Soil type** | **Topography** |
|---|---|---|---|---|---|---|---|---|---|---|
| *Dry zone* | | | | | | | | | | |
| Badenou_1 | Badenou | Dry | 1 | 9.8429 | −5.7094 | 1–10 | Cotton | 1 | Sandy-clayey | Upland |
| Badenou_2 | Badenou | Dry | OGF | 9.8399 | −5.7164 | OGF | — | — | Ferruginous | Slope |
| Badenou_3 | Badenou | Dry | 38 | 9.7814 | −5.6558 | >30 | Rice | 1 | Clayey | Upland |
| Badenou_4 | Badenou | Dry | 12 | 9.7687 | −5.6755 | 11–20 | Maize | 1 | Clayey | Upland |
| Badenou_5 | Badenou | Dry | 21 | 9.7766 | −5.6904 | 21–30 | Maize | 1 | Clayey | Upland |
| Foumbou_1 | Foumbou | Dry | 11 | 8.9136 | −6.0929 | 11–20 | Cotton | 1 | Clayey | Upland |
| Foumbou_2 | Foumbou | Dry | 8 | 8.9089 | −6.1119 | 1–10 | Cotton | 1 | Clayey | Slope |
| Foumbou_3 | Foumbou | Dry | 39 | 8.9084 | −6.1106 | >30 | Yam | 1 | Clayey | Upland |
| Foumbou_4 | Foumbou | Dry | OGF | 8.9179 | −6.0919 | OGF | — | — | Clayey | Upland |
| Foumbou_5 | Foumbou | Dry | 29 | 8.9086 | −6.0956 | 21–30 | Yam | 1 | Ferruginous | Upland |
| *Semi-deciduous zone* | | | | | | | | | | |
| Haut_Sassandra_1 | Haut-Sassandra | Semi-deciduous | 17 | 6.9263 | −6.9040 | 11–20 | Cocoa-coffee | 35 | Ferrallitic | Upland |
| Haut_Sassandra_2 | Haut-Sassandra | Semi-deciduous | 10 | 6.9394 | −6.8863 | 1–10 | Yam | 1 | Ferrallitic | Slope |
| Haut_Sassandra_3 | Haut-Sassandra | Semi-deciduous | 40 | 6.9394 | −6.8746 | >30 | Cocoa-coffee | 40 | Ferrallitic | Slope |
| Haut_Sassandra_4 | Haut-Sassandra | Semi-deciduous | 26 | 6.9433 | −6.8717 | 21–30 | Cocoa | 37 | Ferrallitic | Slope |
| Haut_Sassandra_5 | Haut-Sassandra | Semi-deciduous | OGF | 6.9964 | −6.9135 | OGF | — | — | Ferrallitic | Upland |
| Tene_1 | Téné | Semi-deciduous | 3 | 6.4748 | −5.4016 | 1–10 | Cocoa | 1 | Ferrallitic | Slope |
| Tene_2 | Téné | Semi-deciduous | 21 | 6.4726 | −5.3972 | 21–30 | Coffee | 13 | Ferrallitic | Slope |
| Tene_3 | Téné | Semi-deciduous | OGF | 6.4880 | −5.4976 | OGF | — | — | Ferrallitic | Slope |
| Tene_4 | Téné | Semi-deciduous | 12 | 6.5480 | −5.4391 | 11–20 | Cocoa | 15 | Ferrallitic | Upland |
| Tene_5 | Téné | Semi-deciduous | 43 | 6.5163 | −5.4979 | >30 | Coffee | 1 | Ferrallitic | Slope |
| *Evergreen zone* | | | | | | | | | | |
| Irobo_1 | Irobo | Evergreen | 20 | 5.6155 | −4.7622 | 11–20 | Rice | 1 | Hydromorphic | Lowland |
| Irobo_2 | Irobo | Evergreen | 4 | 5.5513 | −4.7907 | 1–10 | Maize | 1 | Ferrallitic | Upland |
| Irobo_3 | Irobo | Evergreen | OGF | 5.6370 | −4.7478 | OGF | — | — | Ferrallitic | Upland |
| Irobo_4 | Irobo | Evergreen | 23 | 5.6245 | −4.7596 | 21–30 | Maize | 1 | Ferrallitic | Upland |
| Irobo_5 | Irobo | Evergreen | 35 | 5.6429 | −4.7469 | >30 | Rice | 1 | Hydromorphic | Lowland |
| Niegre_1 | Niègré | Evergreen | 10 | 5.1016 | −6.1313 | 1–10 | Cocoa | 1 | Ferrallitic | Slope |
| Niegre_2 | Niègré | Evergreen | OGF | 5.0958 | −6.1579 | OGF | — | — | Ferrallitic | Upland |
| Niegre_3 | Niègré | Evergreen | 19 | 5.0977 | −6.1804 | 11–20 | Cocoa | 36 | Hydromorphic | Lowland |
| Niegre_4 | Niègré | Evergreen | 35 | 5.1236 | −6.1905 | >30 | Cocoa | 40 | Ferrallitic | Slope |
| Niegre_5 | Niègré | Evergreen | 26 | 5.1401 | −6.1881 | 21–30 | Cocoa | 33 | Ferrallitic | Slope |

OGF: old-growth forest reference plot. Age category >30 corresponds to plots aged 30 years or more since agricultural abandonment. Crop year denotes the number of years elapsed since last cultivation at the time of sampling. Dashes (—) indicate that crop type and crop year are not applicable for OGF plots, for which no agricultural legacy is recorded. *Soil types*: ferrallitic = highly weathered tropical soils rich in iron and aluminium oxides; ferruginous = iron-rich soils with moderate weathering; hydromorphic = seasonally waterlogged soils; clayey = fine-textured soils with high clay content; sandy-clayey = intermediate-textured soils with mixed sand and clay fractions. *Topographic positions*: upland = flat or gently sloping interfluve (plateau); slope = inclined terrain (pente); lowland = valley bottom subject to seasonal waterlogging (bas-fond). French vernacular names are given in parentheses to facilitate cross-referencing with local soil and topographic classifications. Phytogeographic zones follow the classification of Guillaumet and Adjanohoun (1971).

### S0. Preliminary analysis: exponential recovery model

#### Model specification

As a first analytical step, we tested whether microbial species richness followed the classical exponential recovery model of Hérault and Piponiot [2018], which assumes a monotonic increase from a post-disturbance baseline $\theta_0$ toward an asymptotic old-growth value $\theta_\infty$:

$$D_{p,t} \sim \text{LogNormal}\big(\log\big[\theta_0 + (\theta_\infty - \theta_0)\left(1 - e^{-\lambda t}\right)\big]\,,\ \sigma_D\big) \tag{1}$$

where $D_{p,t}$ is the species richness of plot $p$ at time $t$ since agricultural abandonment, $\lambda > 0$ is the instantaneous recovery rate, and $\sigma_D$ is the lognormal residual standard deviation. Old-growth forest plots, for which $t$ is undefined, were modelled through a likelihood sharing the asymptotic parameter $\theta_\infty$:

$$D_p \sim \text{LogNormal}(\log \theta_\infty,\ \sigma_{D,\infty}) \tag{2}$$

This formulation allows old-growth plots to directly inform the asymptotic reference value while keeping the recovery trajectory governed by secondary forest plots alone. The model was fitted separately for bacterial and AM fungal richness.

The fundamental biological validity criterion of the exponential recovery model is that $\theta_0 < \theta_\infty$: the post-disturbance baseline must be strictly lower than the old-growth asymptote for the trajectory to describe recovery in the intended direction. Violation of this inequality, i.e. $\theta_0 \geq \theta_\infty$, is diagnostic of model misspecification and indicates that the data are inconsistent with a monotonically increasing recovery toward old-growth levels. The half-life of recovery, defined as the time required to close half the gap between $\theta_0$ and $\theta_\infty$, was derived analytically from posterior samples as:

$$t_{1/2} = \frac{\log 2}{\lambda} \tag{3}$$

#### Results and model rejection

Posterior parameter estimates for both guilds are reported in Table 2. The exponential recovery model failed the primary biological validity criterion for bacterial communities decisively. The posterior mean of the post-disturbance baseline $\theta_0$ (785.2 ASVs; 95% CrI: [570.5, 1129.1]) substantially exceeded the posterior mean of the old-growth asymptote $\theta_\infty$ (724.0 ASVs; 95% CrI: [662.8, 795.1]), with a posterior probability of $\theta_0 > \theta_\infty$ of 1.000 and a posterior mean difference of 746.0 ASVs (95% CrI: [521.0, 1089.1]). This parameter inversion, in which the post-agricultural baseline is estimated with certainty to exceed the old-growth reference, is biologically implausible under the model's assumptions and constitutes unambiguous evidence of misspecification for this guild. It indicates that bacterial richness does not follow a monotonically increasing trajectory toward old-growth levels across the observed chronosequence, consistent with the absence of directional age effects documented in the primary GAM analysis.

For AM fungal communities, the directionality of the model was preserved in the sense that the posterior mean of $\theta_0$ (39.3 taxa; 95% CrI: [12.4, 105.3]) was estimated below $\theta_\infty$ (46.3 taxa; 95% CrI: [33.7, 63.3]), consistent with the expected direction of recovery. However, the credible interval on $\theta_0$ was extremely wide, spanning nearly an order of magnitude and overlapping substantially with the credible interval on $\theta_\infty$. This reflects the high within-category variability in early-successional AM fungal richness visible in Figure 2 and indicates that the model could not reliably distinguish the post-disturbance baseline from the old-growth asymptote given the data. The parametric assumption of a single well-defined post-disturbance starting point is therefore not supported for AM fungi either, even if the directionality constraint is nominally satisfied.

Recovery rates $\lambda$ were similarly estimated for both guilds (AMF: 1.15 [1.02, 1.37]; Bacteria: 1.17 [1.01, 1.40]) and were statistically indistinguishable (P[Bacteria > AMF] = 0.551; posterior mean difference: 0.018; 95% CrI: [-0.250, 0.287]). Derived half-lives were likewise nearly identical between guilds (AMF: 0.60 years [0.51, 0.68]; Bacteria: 0.60 years [0.50, 0.68]; P[Bacteria > AMF] = 0.448; difference: -0.009 [-0.138, 0.124]). Half-life estimates on a sub-annual timescale ($\approx 0.6$ years) for both guilds indicate implausibly rapid estimated early reorganisation. When interpreted in conjunction with the baseline inversion for bacteria and the poorly constrained baseline for AM fungi, these sub-annual half-lives are more parsimoniously explained as artefacts of the exponential model being forced onto data that do not conform to its parametric structure, rather than as genuine biological signals of rapid post-agricultural recovery.

Taken together, these results indicate that the classical exponential recovery model of Hérault and Piponiot [2018] provides an inadequate and biologically misleading description of microbial richness dynamics in this chronosequence, for both guilds and for complementary reasons: parameter inversion in bacteria and unidentifiable baseline in AM fungi.

These failures motivate the adoption of the flexible Bayesian hierarchical GAM framework as the primary analytical approach, which imposes no parametric constraint on the shape of the age trajectory and allows the data to determine whether and in what direction richness changes with successional age.

Table 2: Supplementary Table S0. Posterior parameter estimates from the preliminary exponential recovery model fitted separately to bacterial and AM fungal species richness. Values are posterior means with 95% credible intervals in brackets. $\theta_0$: post-disturbance baseline richness; $\theta_\infty$: old-growth asymptotic richness; $\lambda$: instantaneous recovery rate ($\text{yr}^{-1}$); $\sigma$: lognormal residual standard deviation; $t_{1/2}$: recovery half-life (years). The inversion $\theta_0 > \theta_\infty$ for bacteria (posterior probability = 1.000, highlighted in bold) constitutes unambiguous evidence of model misspecification for this guild. Sub-annual half-life estimates for both guilds further indicate implausibly rapid estimated recovery dynamics inconsistent with the biological interpretation of the model.

| Parameter | Description | AM fungi | Bacteria | P[Bacteria > AMF] |
|---|---|---|---|---|
| $\theta_0$ | Post-disturbance baseline richness | 39.25 [12.43, 105.34] | **785.20 [570.53, 1129.06]** | **1.000** |
| $\theta_\infty$ | Old-growth asymptotic richness | 46.30 [33.73, 63.33] | 724.02 [662.83, 795.13] | 1.000 |
| $\theta_0 > \theta_\infty$ | Inversion probability | — | **1.000** | — |
| $\lambda$ ($\text{yr}^{-1}$) | Instantaneous recovery rate | 1.15 [1.02, 1.37] | 1.17 [1.01, 1.40] | 0.551 |
| $t_{1/2}$ (years) | Recovery half-life | 0.60 [0.51, 0.68] | 0.60 [0.50, 0.68] | 0.448 |
| $\sigma$ | Lognormal residual SD | 0.66 [0.51, 0.86] | 0.20 [0.15, 0.26] | — |
| *Posterior contrasts (Bacteria − AMF)* | | | | |
| $\Delta\theta_0$ | Difference in baseline richness | 746.0 [521.0, 1089.1]; P[Bacteria > AMF] = 1.000 | | |
| $\Delta\theta_\infty$ | Difference in asymptotic richness | 678.0 [612.8, 750.1]; P[Bacteria > AMF] = 1.000 | | |
| $\Delta\lambda$ | Difference in recovery rate | 0.018 [−0.250, 0.287]; P[Bacteria > AMF] = 0.551 | | |
| $\Delta t_{1/2}$ | Difference in half-life | −0.009 [−0.138, 0.124]; P[Bacteria > AMF] = 0.448 | | |

Bold entries indicate the critical model failure: for bacteria, the post-disturbance baseline $\theta_0$ is estimated with posterior probability 1.000 to exceed the old-growth asymptote $\theta_\infty$, violating the fundamental biological assumption of the exponential recovery model. The absence of a guild difference in recovery rate $\lambda$ or half-life $t_{1/2}$ (both $P \approx 0.5$) further indicates that when the exponential model is forced onto these data, it cannot distinguish between guilds in terms of recovery dynamics, consistent with model misspecification rather than genuine biological equivalence.

### S1. Handling of old-growth forest plots in alpha diversity models

Old-growth forest (OGF) plots have no defined successional age, which prevents their direct inclusion in the age smooth $f(\text{age})$. These plots were retained in the model through a binary indicator $1[\text{OGF}]$, which estimates their mean diversity offset from the smooth trajectory evaluated at reference confounder levels. Specifically, the OGF indicator captures the quantity:

$$\beta_{\text{OGF}} = \mathbb{E}[\log \mu_{\text{OGF}}] - f_{\text{guild}}(\overline{\text{age}}_{\text{SF}}) - \beta_{\text{guild}} - \beta_{\text{ref}} \quad (4)$$

where $\overline{\text{age}}_{\text{SF}}$ is the mean secondary forest age used as the smooth evaluation point for OGF predictions, and $\beta_{\text{ref}}$ denotes the fixed effects evaluated at reference confounder levels (crop type = *cacao*, crop year = 0). Former crop type for OGF plots was set to the reference level (*cacao*) and crop year was set to zero, reflecting the absence of agricultural legacy. This formulation treats OGF plots as a structurally distinct category rather than forcing them into the smooth, which would require an arbitrary age assignment and would misrepresent the ecological meaning of the OGF observations.

### S2. Prior specifications

All primary models used weakly informative priors chosen to regularise estimation while remaining diffuse relative to the scale of the data. For fixed regression coefficients the prior was:

$$\beta \sim \mathcal{N}(0,\, 1) \tag{5}$$

on the link scale, which places most prior mass on moderate effect sizes while down-weighting implausibly large effects. Smooth term standard deviations, which control the wiggliness of the spline, received:

$$\sigma_{\text{sds}} \sim \text{LogNormal}(-1,\, 0.5) \tag{6}$$

concentrating mass on smooth rather than highly wiggly trajectories while permitting non-linearity when supported by data. Site-level random effect standard deviations received the same prior:

$$\sigma_{\text{site}} \sim \text{LogNormal}(-1,\, 0.5) \tag{7}$$

Residual standard deviations $\sigma$ for lognormal models also received LogNormal$(-1,\, 0.5)$ priors. The negative binomial shape parameter received a diffuse prior on the positive real line:

$$\phi_{\text{nb}} \sim \text{Gamma}(0.01,\, 0.01) \tag{8}$$

Beta regression precision parameters received:

$$\phi_{\text{beta}} \sim \text{LogNormal}(0,\, 1) \tag{9}$$

All priors were verified through prior predictive checks to ensure they produced plausible diversity values on the response scale before model fitting.

Table 3 summarises the complete prior specification across all model types.

Table 3: Prior distributions used across all Bayesian models. NB = negative binomial; LN = lognormal; Beta = beta regression.

| Parameter | Prior | Models | Interpretation |
|---|---|---|---|
| Fixed coefficients $\beta$ | $\mathcal{N}(0, 1)$ | All | Moderate effects on link scale |
| Smooth SD $\sigma_{\text{sds}}$ | LogNormal$(-1, 0.5)$ | GAM models | Prefers smooth age trajectories |
| Random effect SD $\sigma_{\text{site}}$ | LogNormal$(-1, 0.5)$ | All | Moderate between-site heterogeneity |
| Residual SD $\sigma$ | LogNormal$(-1, 0.5)$ | LN models | Residual variation |
| NB shape $\phi_{\text{nb}}$ | Gamma$(0.01, 0.01)$ | Richness (NB) | Diffuse, positive real line |
| Beta precision $\phi_{\text{beta}}$ | LogNormal$(0, 1)$ | Beta regression | Moderate concentration |

### S3. Marginalisation procedure for posterior predictions

All posterior predictive quantities reported in the main text were marginalised over site random effects by setting `re_formula = NA` in `posterior_epred()`, which integrates over the random effect distribution rather than conditioning on any specific site. The marginalised prediction for a new observation with covariates $\mathbf{x}_*$ and unknown site $s_*$ is:

$$\tilde{\mu}(\mathbf{x}_*) = \int g^{-1}\big(f_{\text{guild}}(\text{age}_*) + \mathbf{x}_*^\top \boldsymbol{\beta} + u_{s_*}\big)\, p(u_{s_*})\, \mathrm{d}u_{s_*} \tag{10}$$

where $g^{-1}(\cdot)$ is the inverse link function and $p(u_{s_*}) = \mathcal{N}(0, \hat{\sigma}^2_{\text{site}})$ uses the posterior mean of the site SD. Confounder variables were fixed at representative values for the secondary forest dataset: the most frequent crop type (*cacao*), the most frequent topography class (*plateau*), the most frequent soil type (*ferralitique*), and the mean crop year (11.5 years). For old-growth forest predictions in the alpha diversity models, crop type was set to the reference level (*cacao*) and crop year to zero. These choices are reported here to ensure reproducibility and to clarify that reported probabilities represent population-level expectations rather than site-specific predictions.

### S4. Confounder treatment in pairwise turnover models

Pairwise dissimilarity data have an inherent non-independence structure: each plot contributes to $n-1$ pairs, meaning observations are not exchangeable. We addressed this through crossed random intercepts:

$$u_i + u_j, \quad u_i,\, u_j \overset{\text{iid}}{\sim} \mathcal{N}(0,\, \sigma^2) \tag{11}$$

where each of the two plots in a pair contributes its own forest-level random intercept. This is a conservative approach that absorbs between-forest baseline differences in composition without assuming independence of pairs sharing a plot.

Pair-level confounder indicators were included as binary fixed effects:

$$c_{ij} = \begin{cases} 1 & \text{if plots } i \text{ and } j \text{ share the same former crop type} \\ 0 & \text{otherwise} \end{cases} \tag{12}$$

$$s_{ij} = \begin{cases} 1 & \text{if plots } i \text{ and } j \text{ share the same soil type} \\ 0 & \text{otherwise} \end{cases} \tag{13}$$

$$t_{ij} = \begin{cases} 1 & \text{if plots } i \text{ and } j \text{ share the same topographic position} \\ 0 & \text{otherwise} \end{cases} \tag{14}$$

These indicators test whether plots sharing land use history or edaphic characteristics are more compositionally similar independently of age, which would confound the age-difference effect $\beta_{\text{age}}$ if unaccounted for.

A sensitivity analysis was conducted by refitting all pairwise turnover models using between-site pairs only (excluding all pairs where $\text{forest}_i = \text{forest}_j$), to assess whether the negative age-difference effects were robust to the potential residual non-independence of within-site pairs. The sensitivity model is:

$$\text{logit}(\delta_{ij}) = \alpha + \beta_{\text{age}}|\text{age}_i - \text{age}_j| + \beta_{\text{crop}}\, c_{ij} + \beta_{\text{soil}}\, s_{ij} + \beta_{\text{topo}}\, t_{ij} + u_i + u_j \tag{15}$$

restricted to pairs where $\text{forest}_i \neq \text{forest}_j$.

### S5. Variance partitioning of compositional variation

To formally assess the relative contributions of successional age, phytogeographic zone, edaphic variables, and site identity to compositional variation in secondary forest plots, we performed a variance partitioning analysis using the `vegan::varpart()` function applied to Hellinger-transformed community matrices. The four predictor sets were:

- $\mathbf{X}_1$: successional age (continuous);
- $\mathbf{X}_2$: phytogeographic zone (categorical, 3 levels: dry, semi-deciduous, evergreen);
- $\mathbf{X}_3$: edaphic variables (soil type and topographic position, categorical);
- $\mathbf{X}_4$: site identity (forest, categorical, 6 levels).

The unique fraction explained by each predictor set, conditioned on all others, is denoted $[a]$ through $[d]$ following the notation of Peres-Neto et al. [2006]:

$$R^2_{\text{adj}}[a] = R^2_{\text{adj}}(\mathbf{X}_1 \mid \mathbf{X}_2, \mathbf{X}_3, \mathbf{X}_4) \tag{16}$$

and analogously for the remaining fractions. We report adjusted $R^2$ values throughout to account for the number of predictors.

**Important caveat.** The six forest sites each belong to a single phytogeographic zone and are characterised by a single dominant soil type and topographic position. This structural design collinearity means that $\mathbf{X}_2$, $\mathbf{X}_3$, and $\mathbf{X}_4$ are not fully separable, and `varpart()` issued collinearity warnings for all multi-table combinations. Individual partitioned fractions should therefore not be interpreted as precise estimates of unique explained variance; they are reported as indicative of the relative importance of each predictor class while acknowledging that shared fractions cannot be cleanly attributed. The total model adjusted $R^2$ and the marginal fractions conditioned on single predictor tables are more reliable than the fully conditioned individual fractions.

### S6. Software and reproducibility

All analyses were conducted in R (v4.5.2; R Core Team 2025). Key packages were:

- `brms` (v2.23.0; Bürkner 2017) for Bayesian model fitting via Stan;
- `vegan` (v2.7.2; Oksanen et al. 2025) for dissimilarity computation, NMDS ordination, PERMANOVA, beta-dispersion analysis, and variance partitioning;
- `tidybayes` (Kay 2024) for posterior extraction and visualisation;
- `ggplot2` (Wickham 2016) and `patchwork` for figure production;
- `purrr` for functional iteration over model combinations;
- `indicspecies` (v1.8; De Cáceres et al. 2010) for indicator species analysis.

Model objects were cached to disk using the `file` argument of `brm()` to ensure reproducibility across sessions. Convergence was verified for all models using the potential scale reduction factor ($\hat{R} < 1.01$ for all parameters) and bulk and tail effective sample size diagnostics ($\mathrm{ESS}_{\mathrm{bulk}} > 400$ and $\mathrm{ESS}_{\mathrm{tail}} > 400$ for all parameters). All code and model output are available in the supplementary data archive.

### Supplementary Table S1. Posterior parameter estimates from Bayesian hierarchical generalised additive models of alpha diversity

Posterior parameter estimates from Bayesian hierarchical generalised additive models of alpha diversity (richness, Shannon, and Simpson) for bacterial and AM fungal communities across a West African forest chronosequence. All parameters are on the log link scale unless otherwise noted. The reference level for crop type is *cacao*; for soil type is *lithosol*; for topography is *sommet*. Guild effect is the contrast of AM fungi relative to bacteria. OGF = old-growth forest. SD = standard deviation of the posterior. 95% CrI = 95% credible interval. All 95% CrIs for confounder levels span zero, indicating no strong evidence of independent edaphic or land-use effects on diversity at the current sample size.

Table 4: Supplementary Table S1. Posterior parameter estimates from Bayesian hierarchical generalised additive models of alpha diversity (richness, Shannon, and Simpson) for bacterial and AM fungal communities across a West African forest chronosequence. All parameters are on the log link scale unless otherwise noted. The reference level for crop type is *cacao*; for soil type is *lithosol*; for topography is *sommet*. Guild effect is the contrast of AM fungi relative to bacteria. OGF = old-growth forest. SD = standard deviation of the posterior. 95% CrI = 95% credible interval. All 95% CrIs for confounder levels span zero, indicating no strong evidence of independent edaphic or land-use effects on diversity at the current sample size. *(Table continues on next page.)*

| Component | Parameter | Metric | Mean | Median | SD | 95% CrI |
|---|---|---|---|---|---|---|
| *Fixed effects* | Intercept (Bacteria, log-scale) | Richness | 6.473 | 6.478 | 0.660 | [5.177, 7.783] |
| | | Shannon | 5.738 | 5.741 | 0.678 | [4.426, 7.054] |
| | | Simpson | 4.962 | 4.962 | 0.711 | [3.546, 6.331] |
| | Guild: AMF vs Bacteria | Richness | −2.711 | −2.711 | 0.089 | [−2.881, −2.533] |
| | | Shannon | −3.485 | −3.488 | 0.128 | [−3.736, −3.233] |
| | | Simpson | −3.125 | −3.128 | 0.156 | [−3.424, −2.812] |
| | Old-growth forest effect | Richness | 0.003 | 0.007 | 0.989 | [−1.987, 1.941] |
| | | Shannon | −0.010 | −0.012 | 1.016 | [−1.973, 1.951] |
| | | Simpson | −0.001 | −0.022 | 1.007 | [−1.960, 1.988] |
| *Crop type*[a] | Cacao-café | Richness | 0.201 | 0.203 | 0.256 | [−0.304, 0.700] |
| | | Shannon | 0.018 | 0.023 | 0.325 | [−0.622, 0.654] |
| | | Simpson | −0.028 | −0.028 | 0.381 | [−0.779, 0.705] |
| | Café | Richness | 0.212 | 0.212 | 0.242 | [−0.274, 0.684] |
| | | Shannon | 0.204 | 0.203 | 0.311 | [−0.415, 0.818] |
| | | Simpson | 0.248 | 0.247 | 0.373 | [−0.478, 0.971] |
| | Coton | Richness | −0.332 | −0.335 | 0.420 | [−1.164, 0.509] |
| | | Shannon | −0.254 | −0.252 | 0.448 | [−1.156, 0.623] |
| | | Simpson | −0.104 | −0.103 | 0.496 | [−1.083, 0.867] |
| | Igname | Richness | 0.325 | 0.322 | 0.469 | [−0.597, 1.283] |
| | | Shannon | 0.661 | 0.672 | 0.519 | [−0.349, 1.679] |
| | | Simpson | 0.813 | 0.818 | 0.571 | [−0.323, 1.920] |
| | Maïs | Richness | −0.094 | −0.092 | 0.334 | [−0.756, 0.570] |
| | | Shannon | −0.005 | 0.001 | 0.376 | [−0.758, 0.725] |
| | | Simpson | −0.069 | −0.063 | 0.408 | [−0.902, 0.717] |
| | Riz | Richness | 0.076 | 0.071 | 0.415 | [−0.733, 0.892] |
| | | Shannon | 0.462 | 0.463 | 0.478 | [−0.506, 1.379] |
| | | Simpson | 0.379 | 0.383 | 0.520 | [−0.646, 1.373] |
| *Crop year*[b] | Years since last cultivation | Richness | −0.001 | 0.000 | 0.007 | [−0.015, 0.013] |
| | | Shannon | 0.002 | 0.002 | 0.009 | [−0.016, 0.019] |
| | | Simpson | 0.000 | 0.000 | 0.011 | [−0.021, 0.022] |

**Supplementary Table S1 (continued).** Posterior parameter estimates from Bayesian hierarchical GAMs of alpha diversity.

| Component | Parameter | Metric | Mean | Median | SD | 95% CrI |
|---|---|---|---|---|---|---|
| *Soil type*[a] | Ferralitique | Richness | 0.036 | 0.032 | 0.347 | [−0.656, 0.731] |
| | | Shannon | 0.142 | 0.149 | 0.379 | [−0.616, 0.889] |
| | | Simpson | 0.220 | 0.226 | 0.422 | [−0.639, 1.021] |
| | Ferrugineux | Richness | 0.016 | 0.015 | 0.289 | [−0.542, 0.597] |
| | | Shannon | 0.242 | 0.243 | 0.384 | [−0.509, 0.988] |
| | | Simpson | 0.296 | 0.295 | 0.467 | [−0.622, 1.214] |
| | Hydromorphe | Richness | −0.160 | −0.162 | 0.641 | [−1.431, 1.103] |
| | | Shannon | 0.244 | 0.237 | 0.663 | [−1.031, 1.565] |
| | | Simpson | 0.462 | 0.460 | 0.684 | [−0.854, 1.828] |
| | Sablo-argileux | Richness | 0.443 | 0.445 | 0.416 | [−0.403, 1.240] |
| | | Shannon | 0.853 | 0.858 | 0.480 | [−0.146, 1.774] |
| | | Simpson | 0.783 | 0.796 | 0.542 | [−0.310, 1.818] |
| *Topography*[a] | Bas-fond | Richness | 0.229 | 0.232 | 0.399 | [−0.562, 0.990] |
| | | Shannon | −0.555 | −0.560 | 0.508 | [−1.536, 0.471] |
| | | Simpson | −0.661 | −0.663 | 0.586 | [−1.797, 0.491] |
| | Pente | Richness | 0.023 | 0.023 | 0.598 | [−1.113, 1.221] |
| | | Shannon | −0.076 | −0.074 | 0.602 | [−1.243, 1.111] |
| | | Simpson | −0.203 | −0.204 | 0.623 | [−1.415, 1.023] |
| | Plateau | Richness | 0.151 | 0.145 | 0.599 | [−1.002, 1.349] |
| | | Shannon | −0.123 | −0.126 | 0.603 | [−1.313, 1.066] |
| | | Simpson | −0.291 | −0.294 | 0.623 | [−1.502, 0.948] |
| *Smooth terms* | $s$(age) — AMF | Richness | 0.325 | 0.296 | 0.153 | [0.120, 0.703] |
| | | Shannon | 0.414 | 0.368 | 0.211 | [0.143, 0.953] |
| | | Simpson | 0.398 | 0.357 | 0.198 | [0.138, 0.888] |
| | $s$(age) — Bacteria | Richness | 0.325 | 0.295 | 0.154 | [0.118, 0.704] |
| | | Shannon | 0.353 | 0.320 | 0.168 | [0.125, 0.771] |
| | | Simpson | 0.361 | 0.327 | 0.175 | [0.127, 0.789] |
| *Variance components* | Site intercept SD | Richness | 0.281 | 0.259 | 0.121 | [0.112, 0.572] |
| | | Shannon | 0.290 | 0.264 | 0.130 | [0.114, 0.608] |
| | | Simpson | 0.301 | 0.275 | 0.134 | [0.115, 0.636] |
| | NB shape $\phi$ | Richness | 14.238 | 13.345 | 5.145 | [6.745, 26.768] |
| | Residual SD $\sigma$ | Shannon | 0.416 | 0.410 | 0.055 | [0.323, 0.541] |
| | | Simpson | 0.526 | 0.520 | 0.069 | [0.410, 0.676] |

[a] Reference levels: crop type = *cacao*; soil type = *lithosol*; topography = *sommet*.
[b] Continuous covariate; effect is per additional year since last cultivation.
All parameters are on the log link scale. Smooth term SDs ($\sigma_{\text{sds}}$) are the standard deviations of the spline basis coefficients and represent the degree of nonlinearity in the estimated age trajectory; larger values indicate more curvature. The negative binomial shape parameter $\phi$ applies to the richness model only; residual SD $\sigma$ applies to the lognormal Shannon and Simpson models. No confounder credible interval excludes zero, consistent with limited statistical power at the current sample size to detect individual edaphic effects.

### Supplementary Table S2. Posterior probability summaries of successional trends

| Metric | Quantity | Estimate |
|---|---|---|
| Richness | $P$(Bacteria increases, age 5 $\rightarrow$ 40) | 0.256 |
| Richness | $P$(AMF increases, age 5 $\rightarrow$ 40) | 0.834 |
| Richness | $P$(AMF trend $>$ Bacteria trend) | 0.772 |
| Richness | $P$(Bacteria at 40 yr $\geq$ OGF level) | 0.482 |
| Richness | $P$(AMF at 40 yr $\geq$ OGF level) | 0.554 |
| Shannon | $P$(Bacteria increases, age 5 $\rightarrow$ 40) | 0.198 |
| Shannon | $P$(AMF increases, age 5 $\rightarrow$ 40) | 0.914 |
| Shannon | $P$(AMF trend $>$ Bacteria trend) | 0.816 |
| Shannon | $P$(Bacteria at 40 yr $\geq$ OGF level) | 0.469 |
| Shannon | $P$(AMF at 40 yr $\geq$ OGF level) | 0.611 |
| Simpson | $P$(Bacteria increases, age 5 $\rightarrow$ 40) | 0.151 |
| Simpson | $P$(AMF increases, age 5 $\rightarrow$ 40) | 0.864 |
| Simpson | $P$(AMF trend $>$ Bacteria trend) | 0.866 |
| Simpson | $P$(Bacteria at 40 yr $\geq$ OGF level) | 0.437 |
| Simpson | $P$(AMF at 40 yr $\geq$ OGF level) | 0.62 |

### Supplementary Table S3. Posterior estimates of the age-difference effect on pairwise compositional dissimilarity

Posterior estimates of $\beta_{\text{age}}$ from Bayesian beta regression models of pairwise compositional dissimilarity, expressing the change in dissimilarity (logit scale) per unit increase in the absolute age difference between plot pairs. Negative values indicate that plots closer in successional age are more compositionally similar. Estimates are marginalised over site random effects and conditioned on pair-level confounders (shared crop type, soil type, and topographic position).

| Guild | Dissimilarity metric | Estimate | 95% CrI |
|---|---|---|---|
| Bacteria | Sørensen | −0.125 | [−0.417, 0.155] |
| Bacteria | Horn | −0.101 | [−0.512, 0.289] |
| Bacteria | Morisita–Horn | −0.110 | [−0.519, 0.275] |
| Bacteria | Bray–Curtis | −0.142 | [−0.456, 0.138] |
| AMF | Sørensen | −0.205 | [−0.537, 0.119] |
| AMF | Horn | −0.322 | [−0.880, 0.163] |
| AMF | Morisita–Horn | −0.317 | [−0.872, 0.157] |
| AMF | Bray–Curtis | −0.279 | [−0.758, 0.107] |

AMF = arbuscular mycorrhizal fungi. Estimates are on the logit scale (i.e. the log-odds of dissimilarity). All 95% CrIs overlap zero, though AMF consistently shows larger negative point estimates than bacteria across all metrics, suggesting a tendency toward convergence in composition with similar successional age. A sensitivity analysis restricted to between-site pairs ($\text{forest}_i \neq \text{forest}_j$) yielded qualitatively identical results (see Section S3).

### Supplementary Figure S1

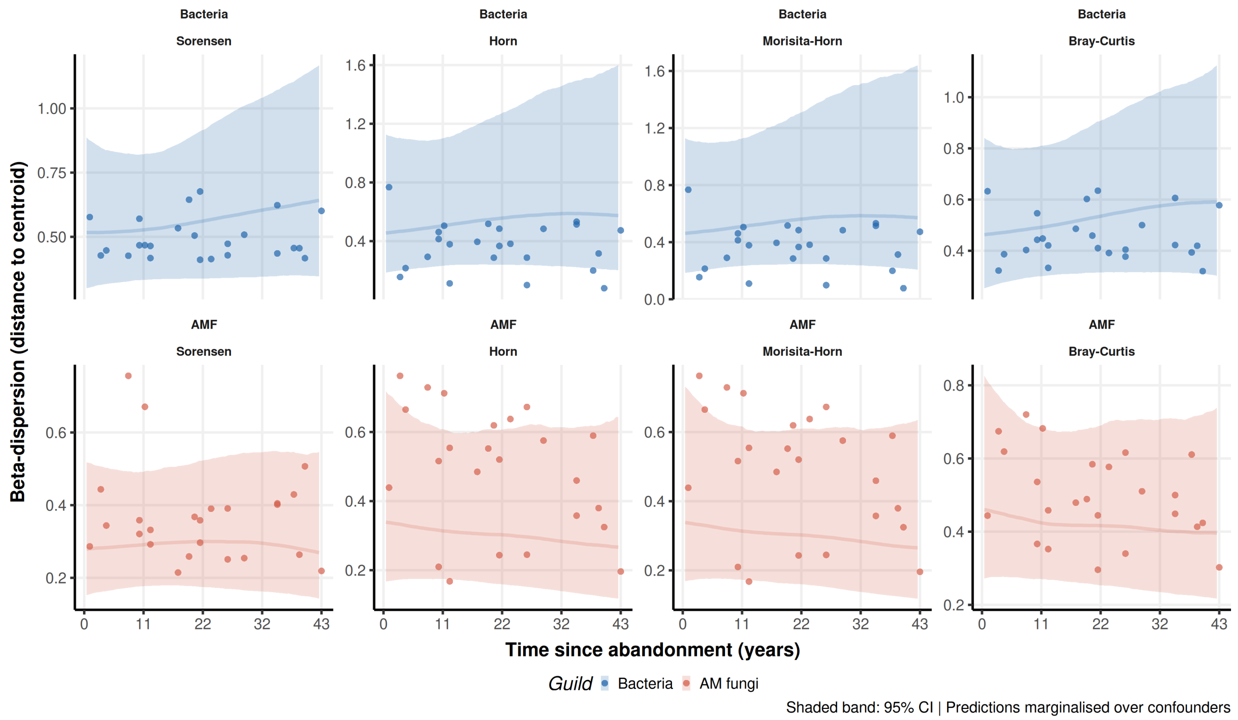


Figure 1: Beta-dispersion of bacterial and AM fungal communities along a West African forest chronosequence. Each panel shows the posterior median predicted distance to forest-level centroid in dissimilarity space (line) and 95% credible interval (shaded band) as a function of time since agricultural abandonment, for four dissimilarity metrics (Sørensen, Horn, Morisita–Horn, Bray–Curtis) and two microbial guilds (bacteria, blue; AM fungi, red). Points represent observed distance-to-centroid values for individual secondary forest plots, computed using `betadisper`. Predictions are marginalised over site random effects and evaluated at the modal land use history and edaphic conditions across the chronosequence. Declining dispersion with age indicates progressive within-forest homogenisation (community stabilisation); increasing or flat dispersion indicates persistent within-forest variability. Bacterial communities show no evidence of declining dispersion across any metric, whereas AM fungal communities show a consistent tendency toward declining dispersion for abundance-weighted metrics (Horn, Morisita–Horn, Bray–Curtis), indicating progressive internal stabilisation of AM fungal composition within forest sites over succession.

### Supplementary Table S4. Posterior summaries of changes in community dispersion

Changes in beta-dispersion (distance to group centroid) from young (5 yr) to old (40 yr) secondary forest, estimated as the difference in posterior mean dispersion between age classes. Positive values indicate increasing dispersion (greater within-group spread) with succession; negative values indicate homogenisation. Estimates are on the dissimilarity scale of each metric. CrI = credible interval.

| Guild | Dissimilarity metric | Estimate | 95% CrI |
|---|---|---|---|
| Bacteria | Sørensen | 0.177 | [−0.125, 0.472] |
| Bacteria | Horn | 0.336 | [−0.266, 0.918] |
| Bacteria | Morisita–Horn | 0.328 | [−0.274, 0.900] |
| Bacteria | Bray–Curtis | 0.195 | [−0.127, 0.515] |
| AMF | Sørensen | −0.011 | [−0.160, 0.146] |
| AMF | Horn | −0.061 | [−0.308, 0.179] |
| AMF | Morisita–Horn | −0.059 | [−0.305, 0.182] |
| AMF | Bray–Curtis | −0.057 | [−0.269, 0.166] |

AMF = arbuscular mycorrhizal fungi. All 95% CrIs span zero, indicating no strong directional trend in community dispersion across succession for either guild. Estimates are derived from Bayesian beta regression models of distance-to-centroid values (betadisper); posterior summaries are marginalised over site random effects.

### Supplementary Table S5. Within-OGF and within-secondary forest beta-dispersion comparison

To assess whether the persistently high compositional distances between secondary forest and old-growth forest (OGF) plots reflect genuine dissimilarity or are partly inflated by heterogeneity among OGF reference plots themselves, we compared mean pairwise distances and beta-dispersion (distance to group centroid) within OGF plots and within secondary forest (SF) plots across all four dissimilarity metrics and both microbial guilds. Results are reported in Table 5.

Table 5: Supplementary Table S5. Comparison of within-group compositional variability between old-growth forest (OGF) and secondary forest (SF) plots across four dissimilarity metrics and two microbial guilds. Mean pairwise distance is the average of all pairwise dissimilarities among plots within each group. Beta-dispersion is the mean distance of each plot to its forest-level group centroid, computed using `betadisper`. $n_{\mathrm{OGF}} = 6$ and $n_{\mathrm{SF}} = 24$ plots across all forests.

| | | Mean pairwise distance | | Mean beta-dispersion | |
|---|---|---|---|---|---|
| Guild | Metric | OGF | SF | OGF | SF |
| Bacteria | Sørensen | 0.900 | 0.866 | —[a] | 0.478 |
| | Horn | 0.818 | 0.728 | —[a] | 0.352 |
| | Morisita–Horn | 0.817 | 0.727 | —[a] | 0.352 |
| | Bray–Curtis | 0.877 | 0.823 | —[a] | 0.442 |
| AM fungi | Sørensen | 0.697 | 0.603 | —[a] | 0.356 |
| | Horn | 0.891 | 0.769 | —[a] | 0.462 |
| | Morisita–Horn | 0.891 | 0.769 | —[a] | 0.462 |
| | Bray–Curtis | 0.865 | 0.782 | —[a] | 0.478 |

[a] OGF beta-dispersion could not be computed because each forest site contributes a single OGF plot, making each plot its own group centroid and yielding a distance of zero by definition. Mean pairwise distance across OGF plots therefore provides the only available measure of between-OGF compositional variability, and should be interpreted as a landscape-scale index of OGF heterogeneity rather than within-site dispersion.

Within-OGF mean pairwise distances are consistently high across all metrics and both guilds (range: 0.697–0.900), comparable to or exceeding within-SF mean pairwise distances (range: 0.603–0.866), indicating that OGF plots are themselves compositionally heterogeneous at the landscape scale. By contrast, within-SF beta-dispersion is substantially larger than within-OGF dispersion at the site level, reflecting the greater compositional variability among secondary forest plots within individual forest sites relative to their OGF references. These results indicate that the persistently high compositional distances between secondary and old-growth forest communities primarily reflect genuine dissimilarity rather than artefactual inflation by OGF heterogeneity, though cross-site variability among OGF communities introduces uncertainty into landscape-level recovery assessments that cannot be resolved with the current reference plot density of one plot per forest site.

### Supplementary Table S6. Variance partitioning of compositional variation

Variance partitioning was performed on Hellinger-transformed community matrices for secondary forest plots ($n = 24$) to assess the relative contributions of successional age ($\mathbf{X}_1$), phytogeographic zone ($\mathbf{X}_2$), edaphic variables ($\mathbf{X}_3$), and site identity ($\mathbf{X}_4$) to bacterial and AM fungal compositional variation. Results are reported in Tables 6 and 7. The significance of the unique age fraction, conditioned on all other predictors, was assessed via partial redundancy analysis with 9,999 permutations (Table 8).

Strong collinearity was detected among predictor sets $\mathbf{X}_2$, $\mathbf{X}_3$, and $\mathbf{X}_4$ in all multi-table combinations, reflecting the structural design constraint that each forest site belongs to a single phytogeographic zone and is characterised by a single dominant soil type and topographic position. As a consequence, individual partitioned fractions involving these predictors jointly should not be interpreted as precise estimates of unique explained variance; they are reported here for completeness. The most reliable quantities are the total model adjusted $R^2$, the unique fraction of age conditioned on all other predictors ($[a] = \mathbf{X}_1 \mid \mathbf{X}_2, \mathbf{X}_3, \mathbf{X}_4$), and the marginal fractions conditioned on single predictor tables.

#### Key results

For bacterial communities, the full model explained 20.8% of compositional variation (total Adj. $R^2 = 0.208$). Successional age contributed a negligible unique fraction when conditioned on all other predictors ($[a] = 0.003$), while site identity ($[d] = 0.077$ conditioned on zone, edaphics, and age) and edaphic variables ($[c] = 0.032$) explained larger independent fractions. Partial RDA confirmed that the unique age effect was non-significant after controlling for all other predictors ($F_{1,12} = 1.04$, $P = 0.329$).

For AM fungal communities, the full model explained only 3.9% of compositional variation (total Adj. $R^2 = 0.039$), with the majority of variation unexplained ($[p] = 0.961$). Successional age contributed near-zero or negative unique fractions under all conditioning combinations ($[a] = 0.000$ conditioned on all others), and the partial RDA was likewise non-significant ($F_{1,12} = 1.00$, $P = 0.434$). Edaphic variables showed the largest unique fraction ($[c] = 0.049$), though this too was subject to collinearity caveats.

Taken together, these results confirm that successional age does not exert a detectable independent effect on community composition in either guild after accounting for site-level environmental heterogeneity, and that the compositional signal attributed to age in ordination space is substantially confounded with phytogeographic zone and site identity.

Table 6: Supplementary Table S4a. Variance partitioning of Hellinger-transformed bacterial community composition for secondary forest plots ($n = 24$). Adjusted $R^2$ values are reported throughout. Predictor sets: $\mathbf{X}_1$ = successional age; $\mathbf{X}_2$ = phytogeographic zone; $\mathbf{X}_3$ = edaphic variables (soil type, topography); $\mathbf{X}_4$ = site identity. Total variation SS = 17.203; total variance = 0.748. Individual fractions [$e$]–[$o$] represent shared and joint components; only testable unique fractions are highlighted.

| Fraction | Description | Adj. $R^2$ | Testable |
|---|---|---|---|
| *Marginal fractions (single predictors)* | | | |
| $[\mathbf{X}_1]$ | Successional age alone | 0.002 | Yes |
| $[\mathbf{X}_2]$ | Phytogeographic zone alone | 0.102 | Yes |
| $[\mathbf{X}_3]$ | Edaphic variables alone | 0.107 | Yes |
| $[\mathbf{X}_4]$ | Site identity alone | 0.170 | Yes |
| *Combined fractions* | | | |
| $[\mathbf{X}_1 + \mathbf{X}_2]$ | Age + zone | 0.108 | Yes |
| $[\mathbf{X}_1 + \mathbf{X}_3]$ | Age + edaphics | 0.109 | Yes |
| $[\mathbf{X}_1 + \mathbf{X}_4]$ | Age + site | 0.176 | Yes |
| $[\mathbf{X}_2 + \mathbf{X}_3]$ | Zone + edaphics | 0.128 | Yes |
| $[\mathbf{X}_2 + \mathbf{X}_4]$ | Zone + site | 0.170 | Yes |
| $[\mathbf{X}_3 + \mathbf{X}_4]$ | Edaphics + site | 0.205 | Yes |
| [All] | Full model | **0.208** | Yes |
| *Unique fractions (conditioned on all others)* | | | |
| $[a] = \mathbf{X}_1 \mid \mathbf{X}_2, \mathbf{X}_3, \mathbf{X}_4$ | Unique age | 0.003 | Yes |
| $[b] = \mathbf{X}_2 \mid \mathbf{X}_1, \mathbf{X}_3, \mathbf{X}_4$ | Unique zone | 0.000 | Yes |
| $[c] = \mathbf{X}_3 \mid \mathbf{X}_1, \mathbf{X}_2, \mathbf{X}_4$ | Unique edaphics | 0.032 | Yes |
| $[d] = \mathbf{X}_4 \mid \mathbf{X}_1, \mathbf{X}_2, \mathbf{X}_3$ | Unique site | 0.077 | Yes |
| $[p]$ = Residuals | Unexplained | 0.792 | — |

Collinearity was detected among $\mathbf{X}_2$, $\mathbf{X}_3$, and $\mathbf{X}_4$ in all multi-table combinations. Individual partitioned fractions involving these predictors jointly may be unreliable; the unique fractions [$a$]–[$d$] and the full-model $R^2$ are the most interpretable quantities.

Table 7: Supplementary Table S4b. Variance partitioning of Hellinger-transformed AM fungal community composition for secondary forest plots ($n = 24$). Layout and notation as in Table S4a. Total variation SS = 14.471; total variance = 0.629. The near-zero total model adjusted $R^2$ and dominant residual fraction indicate that the four predictor sets collectively explain very little AM fungal compositional variation in this dataset.

| Fraction | Description | Adj. $R^2$ | Testable |
|---|---|---|---|
| *Marginal fractions (single predictors)* | | | |
| $[\mathbf{X}_1]$ | Successional age alone | −0.007 | Yes |
| $[\mathbf{X}_2]$ | Phytogeographic zone alone | 0.002 | Yes |
| $[\mathbf{X}_3]$ | Edaphic variables alone | 0.043 | Yes |
| $[\mathbf{X}_4]$ | Site identity alone | −0.002 | Yes |
| *Combined fractions* | | | |
| $[\mathbf{X}_1 + \mathbf{X}_2]$ | Age + zone | −0.006 | Yes |
| $[\mathbf{X}_1 + \mathbf{X}_3]$ | Age + edaphics | 0.043 | Yes |
| $[\mathbf{X}_1 + \mathbf{X}_4]$ | Age + site | −0.009 | Yes |
| $[\mathbf{X}_2 + \mathbf{X}_3]$ | Zone + edaphics | 0.029 | Yes |
| $[\mathbf{X}_2 + \mathbf{X}_4]$ | Zone + site | −0.002 | Yes |
| $[\mathbf{X}_3 + \mathbf{X}_4]$ | Edaphics + site | 0.039 | Yes |
| [All] | Full model | **0.039** | Yes |
| *Unique fractions (conditioned on all others)* | | | |
| $[a] = \mathbf{X}_1 \mid \mathbf{X}_2, \mathbf{X}_3, \mathbf{X}_4$ | Unique age | 0.000 | Yes |
| $[b] = \mathbf{X}_2 \mid \mathbf{X}_1, \mathbf{X}_3, \mathbf{X}_4$ | Unique zone | 0.000 | Yes |
| $[c] = \mathbf{X}_3 \mid \mathbf{X}_1, \mathbf{X}_2, \mathbf{X}_4$ | Unique edaphics | 0.049 | Yes |
| $[d] = \mathbf{X}_4 \mid \mathbf{X}_1, \mathbf{X}_2, \mathbf{X}_3$ | Unique site | 0.011 | Yes |
| $[p]$ = Residuals | Unexplained | 0.961 | — |

Collinearity warnings apply as in Table S4a. The dominant residual fraction (0.961) indicates that the measured predictors collectively account for very little AM fungal compositional variation, consistent with strong stochastic assembly and high unexplained site-level heterogeneity.

Table 8: Supplementary Table S4c. Partial redundancy analysis tests for the significance of the unique successional age fraction in bacterial and AM fungal community composition, conditioned on phytogeographic zone, edaphic variables, and site identity. Permutation tests were performed with 9,999 free permutations.

| Guild | Df (model) | Df (residual) | $F$ | $P$ |
|---|---|---|---|---|
| Bacteria | 1 | 12 | 1.043 | 0.329 |
| AM fungi | 1 | 12 | 1.003 | 0.434 |

Neither guild shows a statistically significant unique age effect after conditioning on zone, edaphic variables, and site identity, confirming that successional age does not independently structure community composition beyond what is explained by site-level environmental heterogeneity in this dataset.

Jari Oksanen, Gavin L. Simpson, F. Guillaume Blanchet, Roeland Kindt, Pierre Legendre, Peter R. Minchin, R. B. O'Hara, Peter Solymos, M. Henry H. Stevens, Eduard Szoecs, Helene Wagner, Matt Barbour, Michael Bedward, Ben Bolker, Daniel Borcard, Tuomas Borman, Gustavo Carvalho, Michael Chirico, Miquel De Caceres, Sebastien Durand, Heloisa Beatriz Antoniazi Evangelista, Rich FitzJohn, Michael Friendly, Brendan Furneaux, Geoffrey Hannigan, Mark O. Hill, Leo Lahti, Dan McGlinn, Marie-Helene Ouellette, Eduardo Ribeiro Cunha, Tyler Smith, Adrian Stier, Cajo J. F. Ter Braak, and James Weedon. *Vegan: Community Ecology Package*. 2025.

Pedro R. Peres-Neto, Pierre Legendre, Stéphane Dray, and Daniel Borcard. Variation partitioning of species data matrices: Estimation and comparison of fractions. *Ecology*, 87(10):2614–2625, October 2006. ISSN 0012-9658. doi: 10.1890/0012-9658(2006)87[2614:vposdm]2.0.co;2.

R Core Team. *R: A Language and Environment for Statistical Computing*. R Foundation for Statistical Computing, Vienna, Austria, 2025.

Hadley Wickham. *ggplot2: Elegant Graphics for Data Analysis*. Springer-Verlag New York, 2016. ISBN 978-3-319-24277-4. URL `https://ggplot2.tidyverse.org`.
